# Endothelial cannabinoid CB1 receptor deficiency reduces arterial inflammation and lipid uptake in response to atheroprone shear stress

**DOI:** 10.1101/2024.05.15.594375

**Authors:** Bingni Chen, Aishvaryaa Prabhu, Guo Li, Anna Kaltenbach, Yong Wang, George Shakir, Lucia Natarelli, Remco Megens, Yvonne Jansen, Srishti Ramanathan, Martina Geiger, Alexander Faussner, Michael Hristov, Daniel Richter, Xinyu Di, Mario van der Stelt, Zhaolong Li, Nadja Sachs, Valentina Paloschi, Lars Maegdefessel, Susanna M. Hofmann, Christian Weber, Stephan Herzig, Raquel Guillamat Prats, Sabine Steffens

## Abstract

**Background:** Peripherally restricted cannabinoid CB1 receptor antagonists without central side effects hold promise for treating metabolic disorders including diabetes and obesity. In atherosclerosis, the specific effects of peripheral CB1 signaling in vascular endothelial cells (ECs) remain incompletely understood.

**Methods and results:** Endothelial expression of the CB1 encoding gene *CNR1* was detectable in human plaque single-cell RNA sequencing data. *In situ* hybridization of *Cnr1* in murine aortas revealed a significantly increased endothelial expression within atheroprone compared to atheroresistant regions. *In vitro*, *CNR1* was upregulated by oscillatory shear stress (OSS) in human aortic endothelial cells (HAoECs). Endothelial CB1 deficiency (*Cnr1*^EC-KO^) in female mice on atherogenic background resulted in pronounced endothelial phenotypic changes, with reduced vascular inflammation and permeability. This resulted in attenuated plaque development with reduced lipid content in female mice, while reduced white and brown adipose tissue mass and liver steatosis were observed in both males and females. *Ex vivo* imaging of carotid arteries via two-photon microscopy revealed less labeled low density lipoprotein (Dil-LDL) uptake in *Cnr1*^EC-KO^. This was accompanied by a significant reduction of aortic endothelial caveolin-1 (CAV1) expression, a key structural protein involved in lipid transcytosis, in female *Cnr1*^EC-KO^ mice. *In vitro*, pharmacological blocking with CB1 antagonist AM281 reproduced the inhibition of CAV1 expression and Dil-LDL uptake in response to atheroprone OSS in HAoECs, which was dependent on cAMP-mediated PKA activation. Conversely, the CB1 agonist ACEA increased Dil-LDL uptake and CAV1 expression in HAoECs. Finally, treatment of atherosclerotic mice with the peripheral CB1 antagonist JD-5037 reduced plaque progression, CAV1 and endothelial adhesion molecule expression in female mice.

**Conclusions:** These results confirm an essential role of endothelial CB1 to the pathogenesis of atherosclerosis. Peripheral CB1 antagonists may hold promise as an effective therapeutic strategy for treating atherosclerosis and related metabolic disorders.

**NOVELTY AND SIGNIFICANCE:** **What is known?**

- Enhanced endocannabinoid-cannabinoid CB1 receptor signaling has been implicated in metabolic disorders, atherosclerosis, and hypertension, but the cell-specific role of endothelial CB1 in atherosclerosis is not well understood.
- Global CB1 antagonists improve metabolic function and inhibit atherosclerotic plaque development in mouse models, but have failed in the clinic due to centrally mediated psychiatric side effects.

**What is new?**

- Based on human single-cell RNA sequencing data, CB1 is expressed in human plaque ECs.
- Using transgenic mouse models and human primary aortic endothelial cells, we provide evidence for a key role of CB1 in endothelial shear stress response, inflammatory gene expression, and LDL uptake.
- The underlying signaling pathway of CB1-induced endothelial LDL uptake involves a cAMP-PKA-dependent regulation of caveolin 1 (CAV1) expression, a structural protein of the shear stress sensitive signaling domains of the plasma membrane.
- By limiting endothelial CAV1 and VCAM1 expression, peripherally restricted CB1 antagonists confer atheroprotection in mice.

Our findings reveal that endothelial CB1 expression is induced by atheroprone shear stress responses and contributes to impaired vascular barrier function, inflammation, and lipid uptake, thereby promoting atherosclerotic lesion formation and progression. By elucidating the transcriptomic pathways regulated by endothelial CB1 and its broad influence on lipid uptake and metabolism in arteries, liver, and brown and white adipose tissue, our study provides insights into novel pathways and potential interventions for the treatment of atherosclerosis and metabolic disorders. The use of peripherally restricted CB1 antagonists that specifically target vascular inflammation and tissue lipid storage could be a complementary and safe therapeutic avenue to treat cardiovascular and metabolic disease comorbidities without altering CB1 signaling in the brain.

## INTRODUCTION

Atherosclerosis is a lipid-driven, pro-inflammatory autoimmune disease characterized by the progressive accumulation of lipids, inflammatory cells, and fibrous material in the subendothelial space of large arteries, leading to life-threatening complications, including myocardial infarction, stroke, and peripheral artery disease.^1^ Accumulation of LDL in the arterial wall and endothelial dysfunction are thought to be key mechanisms that initiate the disease.^2^ Atherosclerotic lesions develop preferentially at arterial branches or curvatures of blood vessels, where blood flow is disturbed, further highlighting the importance of biomechanical factors in the initiation of the disease.^3^ Shear stress, the frictional force generated by the blood flow on the endothelial surface, is sensed by mechanoreceptors and induces an intracellular signaling response. In atheroprone regions with disturbed flow, the endothelium is exposed to low, oscillatory shear stress, which induces inflammatory signaling through activation of the nuclear factor-kappa-B (NF-κB) pathway. In contrast, atheroprotective shear stress induces several negative regulators of inflammatory pathways including the transcription factor Kruppel-like family 2 (KLF2).^4^ As a result, in regions of atheroprotective shear stress the endothelium has a quiescent phenotype with a preserved barrier function, whereas low, oscillatory shear stress leads to an activated phenotype with a relocalization of intercellular junctional complexes and increased permeability.^2,3^

A key regulator linking biomechanical and inflammatory pathways to LDL transport into the arterial wall is endothelial caveolin 1 (CAV1), the major component of caveolae.^5^ Caveolae are bulb-shaped, shear stress-sensitive signaling domains in the plasma membrane of endothelial cells.^5^ Genetic loss of *Cav-1* reduced LDL infiltration into the arterial wall, promoted nitric oxide production, and reduced the expression of adhesion molecules, while all these effects were reversed by endothelial-specific *Cav1* transgene expression.^6^ CAV1 is required for caveolae formation and caveolae-mediated LDL uptake and transcytosis across the endothelium. LDL transcytosis pathways involve scavenger receptor B1 (SR-B1) and activin-like kinase 1 (ALK1), which are localized within caveolae and directly bind LDL to facilitate the transport of intact LDL across endothelial cells.^7,8^ This is independent of the canonical LDLR-dependent LDL endocytosis pathway with lysosomal degradation and instead mediates LDL translocation across the endothelial layer to become trapped in the vessel wall.^5^

Endocannabinoids, a group of arachidonic acid-derived lipid mediators that bind to the primary G protein-coupled receptors CB1 and CB2, play an important role in energy homeostasis and metabolic health.^9^ In obese patients, elevated endocannabinoid levels have been positively correlated with coronary endothelial dysfunction.^10^ Pharmacologic blockade of CB1 signaling with the antagonist rimonabant inhibited plaque formation in a mouse model of atherosclerosis.^11^ Other investigators did not find an effect on the plaque size, but observed an improved aortic endothelium-dependent vasodilation and decreased aortic ROS production and NADPH oxidase activity in mice treated with CB1 antagonist.^12^ However, the beneficial use of global CB1 antagonists has been hampered by serious side effects, particularly depression and anxiety, due to the inhibition of CB1 signaling in the brain.

Peripherally restricted CB1 antagonists have been developed and shown to ameliorate metabolic dysfunction in mouse models.^13,14^ Furthermore, adipocyte-specific deletion of the CB1 encoding gene *Cnr1* was sufficient to protect mice from diet-induced obesity and associated metabolic alterations.^15^ We have recently shown that myeloid-specific *Cnr1* deficiency limits atherosclerosis by inhibiting the recruitment and proliferation of arterial macrophages.^16^ However, the specific role of endothelial CB1 in atherogenesis and related metabolic alterations remains poorly defined. Therefore, to elucidate the endothelial role of CB1 in atherosclerosis, we generated an endothelial-specific *Cnr1* knockout mouse line and provide evidence for a key role of CB1 in endothelial shear stress response, inflammatory gene expression, and LDL uptake. These findings were supported by mechanistic *in vitro* experiments with primary HAoECs and *in vivo* experiments with a peripheral antagonist, indicating that peripheral CB1 antagonists may hold promise as an effective therapeutic strategy for treating atherosclerosis and related metabolic disorders.

## METHODS

### Data Availability

A detailed description of the materials and methods is provided in the Supplemental Material. The RNA sequencing data is accessible under GEO accession number GSE260826. Additional data supporting the findings of this study are available from the corresponding authors on request.

## RESULTS

### Endothelial CB1 expression is induced by atheroprone shear stress

While CB1 expression has been reported in human plaques,^17^ the specific cell types contributing to CB1 signaling within the plaque have not been well characterized. Therefore, we screened single-cell RNA sequencing data from human carotid plaques^18^ and found that the CB1-encoding gene *CNR1* was detectable in plaque endothelial cells (Figure 1A-B). *CNR1* expression was not detectable in other cell clusters of human atherosclerotic plaques identified by single cell RNA sequencing, although the expression has previously been demonstrated in a variety of immune cells^19^ and more recently in murine plaque macrophages by *in situ* hybridization.^16^

**Figure 1.**
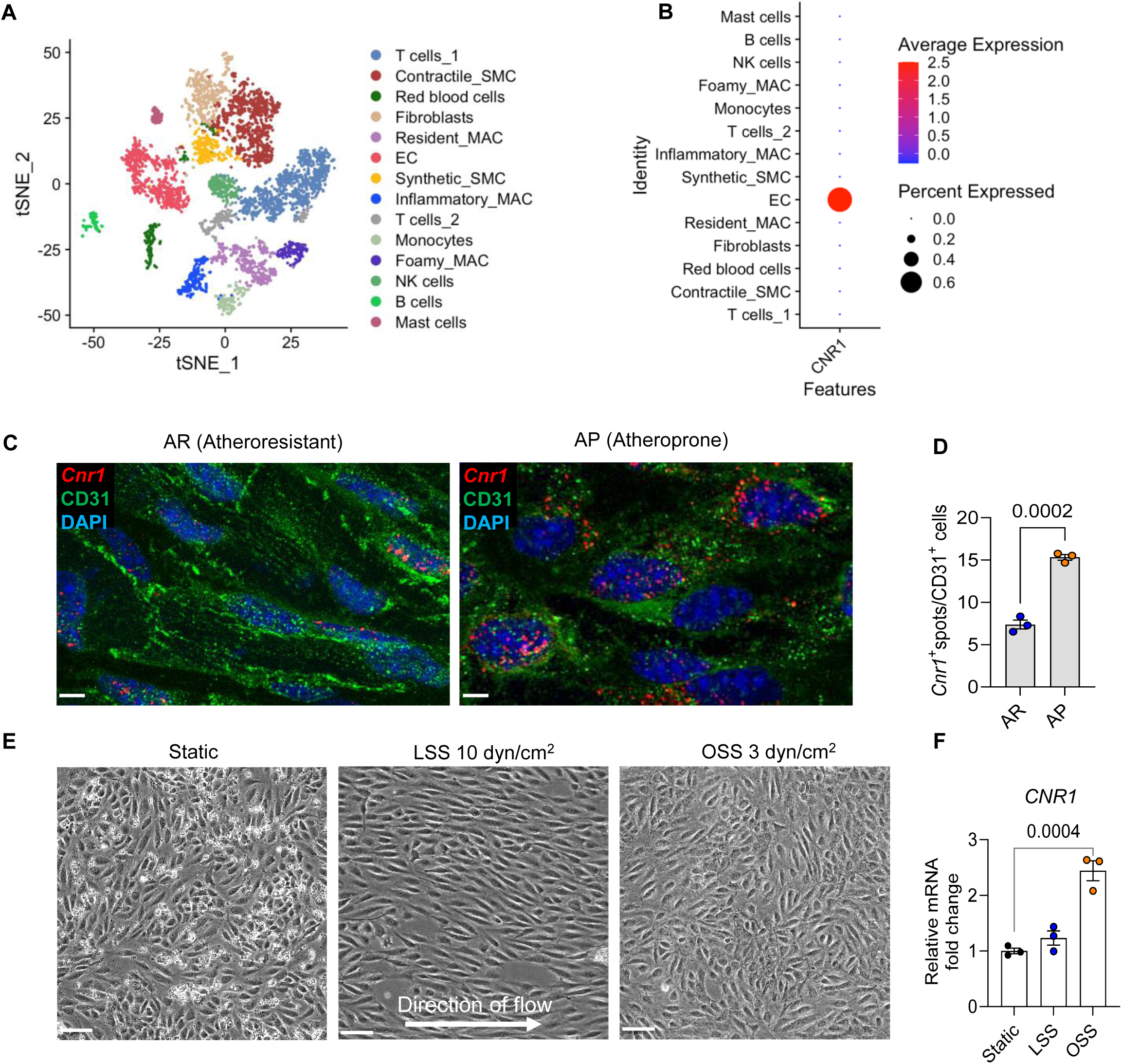
Impact of shear stress on endothelial CB1 expression. (**A**) t-distributed stochastic neighbor embedding (t-SNE) reveals cell clusters in human atherosclerosis single-cell RNA sequencing (scSeq) data provided by the Munich Vascular Biobank. (**B**) The dot plot displays the *CNR1* expression patterns within each cell cluster, with the colour gradient from blue to red indicating the average expression level (low to high) and the point size reflecting the percentage of gene expression within each cluster. (SMC: smooth muscle cells, MAC: macrophage, EC: endothelial cells, NK cell: natural killer cells) (**C**) En face *in situ* hybridization for *Cnr1* within the descending thoracic aorta (atheroresistant, AR) and aortic arch (atheroprone, AP) of the same *Apoe^−/−^* mouse. Scale bar, 5 μm. (**D**) A total of 3 *Apoe^−/−^* mice were included in the analysis; 10-15 images per mouse were acquired for quantification. (**E**) Representative images of HAoECs after 24 h exposure to LSS (10 dyn/cm^2^) or OSS (3 dyn/cm^2^) compared to static conditions. Scale bar, 50 μm. (**F**) *CNR1* gene expression levels in HAoECs after exposure to different shear stress (n=3 independent experiments). Data are shown as mean ± s.e.m.; (**D**) unpaired Student t tests and (**F**) one-way ANOVA followed by a post-hoc Tukey multi-comparison test were used to determine the significant differences.

ECs exhibit distinct phenotypes instructed by their specific location within the artery. In the descending thoracic aorta, the endothelium is exposed to a uniform and laminar blood flow, whereas within the inner curvature of the aortic arch or at branch points, the blood flow is disturbed.^20^ We investigated whether the expression of endothelial CB1 is affected by shear stress. To this end, we first performed an *en face in situ* hybridization to detect the mRNA expression of endothelial *Cnr1* in different shear stress regions within the aorta of 8-week-old *Apoe^−/−^* mice. A more intense endothelial *Cnr1* expression signal was detected in the atheroprone region of the aortic arch, compared to the atheroresistant region of the descending thoracic aorta (Figure 1C-D, Supplemental Figure S1A-B). This observation suggests that the expression of endothelial *Cnr1* is regulated by shear stress.

To further demonstrate this regulation, we cultured HAoECs in different shear stress conditions. After 24 h of exposure to laminar shear stress (LSS), HAoECs had a more elongated shape and were aligned in flow direction as opposed to cells cultured in static conditions, whereas OSS resulted in a cobblestone-like morphology (Figure 1E). In line with the *in vivo* findings in *Apoe^−/−^* aortas, 2-fold higher *CNR1* mRNA levels were observed in HAoECs exposed to OSS compared to static culture or LSS, while LSS did not affect *CNR1* expression (Figure 1F). No significant differences in the mRNA levels of the endocannabinoid 2-AG synthesis enzyme *DAGL* and degradation enzyme *MAGL* were detectable in response to different shear stress conditions (Supplemental Figure S1C). Collectively, these data indicate that endothelial CB1 expression is induced by OSS in atheroprone regions of the aorta.

### Endothelial *Cnr1* deficiency affects vascular and cardiac function

Next, we generated endothelial cell-specific *Cnr1* deficient mice by crossing *Cnr1^flox/flox^* mice^21^ with *Bmx^CreERT^* mice^22^ on an *Apoe^−/−^* background (*Apoe^−/−^Bmx^Cre(+/-)^Cnr1^flox/flox^*), hereafter referred to as *Cnr1^EC-KO^* mice. Tamoxifen-induced recombination at 8 weeks of age resulted in arterial endothelial cell-specific deletion of *Cnr1* (Supplemental Figure S2A).^22^ Tamoxifen was also administered to the control group (*Apoe^−/−^Bmx^Cre(+/-)^* mice, designated *Cnr1^EC-WT^*). *Cnr1* deletion was confirmed by *in situ* hybridization in *Cnr1^EC-KO^*aortic roots, and the endothelial CB1 expression in *Cnr1^EC-WT^*mice was confirmed by colocalization of the *Cnr1* probe with the endothelial marker CD31 (Supplemental Figure S2B).

To perform a basic cardiovascular phenotyping of the new mouse line, we measured cardiac function and blood flow dynamics in *Cnr1^EC-KO^* and *Cnr1^EC-WT^* mice by echocardiography. While *Cnr1^EC-WT^* mice showed impaired cardiac function after 4 weeks of Western diet (WD), characterized by reduced fractional shortening and ejection fraction, these parameters were preserved in *Cnr1^EC-KO^* mice (Supplemental Figure S3A-3D). Notably, we observed an increase in flow velocity in atheroprone regions but not the atheroresistant region of the aorta in *Cnr1^EC-KO^* mice compared to *Cnr1^EC-WT^*mice (Supplemental Figure S3E-G).

### Endothelial *Cnr1* deficiency affects aortic endothelial morphology and integrity

Given that OSS induces endothelial CB1 expression, we asked whether the absence of endothelial CB1 would in turn affect the endothelial phenotype in response to OSS. Therefore, we performed *en face* staining for the EC junctional protein vascular endothelial (VE)-cadherin and the adhesion molecule ICAM1 using aortic arches and thoracic aortas isolated from *Cnr1^EC-KO^*and *Cnr1^EC-WT^* mice. Increased VE-cadherin expression was observed in the atheroresistant descending thoracic aorta of *Cnr1^EC-KO^* and *Cnr1^EC-WT^* mice (Figure 2A-B). ECs in *Cnr1^EC-KO^* aortas were more elongated than those in *Cnr1^EC-WT^* aortas (Figure 2C). Furthermore, less endothelial ICAM1 expression was found in aortas of *Cnr1^EC-KO^* mice, suggesting an anti-inflammatory phenotype when endothelial *Cnr1* is depleted (Figure 2D). Because vascular inflammation impairs endothelial barrier function, we subsequently monitored vascular leakage by injecting Evańs Blue into *Cnr1^EC-KO^* and *Cnr1^EC-WT^* mice after 4 weeks of WD. Significantly less vascular leakage was observed in the aortas of *Cnr1^EC-KO^* mice, particularly at atheroprone sites such as the inner curvature of the aortic arch, indicating preserved endothelial integrity in *Cnr1^EC-KO^* mice (Figure 2E-F).

**Figure 2.**
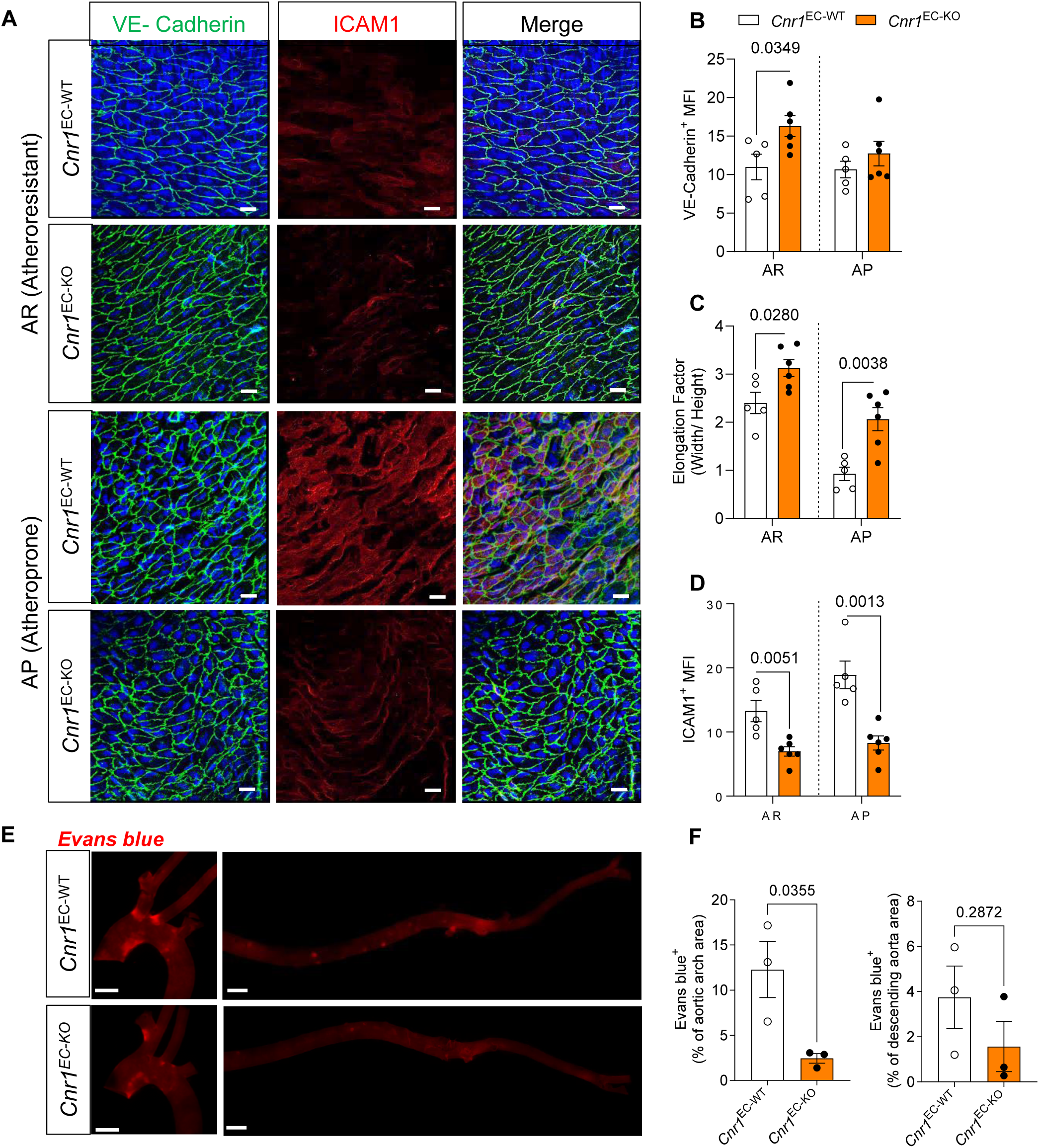
Impact of endothelial *Cnr1* deficiency on endothelial morphology and integrity. (**A**) The aortic arch (atheroprone, AR) and descending thoracic aorta (atheroresistant, AR) were isolated from female *Cnr1^EC-WT^* mice (n = 5) and *Cnr1^EC-KO^* mice (n = 6) and prepared for en face staining. Staining for VE-cadherin (green), ICAM1 (red), and DAPI (blue) to visualize the intercellular junctions, adhesion molecules and nuclei. The images were captured by confocal microscopy. Scale bar, 20 μm. (**B-D**) The mean fluorescence intensity (MFI) of VE-cadherin and ICAM1 signals were quantified from maximum projections of images generated from Z-stack (10-15 images per region per mouse were captured) with Leica software. EC elongation factor (of 20 randomly selected cells per image) was quantified with Image J. (**E**) Representative images of endothelial permeability assay, based on Evańs blue extravasation into aortas 30 min after i.v. injection. Scale bar, 1 mm. (**F**) Quantification of Evańs blue positive area normalized to aortic arch or descending aorta area of female *Cnr1^EC-WT^* and *Cnr1^EC-KO^* mice (n =3). Images were taken with a Leica DM6000B microscope and quantified by LAS V4.3 software. Data are shown as mean ± s.e.m. and unpaired Student’s t-test was performed; AR and AP data shown in **B-D** were analysed separately.

### Transcriptomic profiling of *Cnr1^EC-KO^*aortic endothelial cells reveals profound transcriptomic changes

To globally assess the transcriptomic profile regulated by endothelial CB1 signaling, we sorted aortic ECs from *Cnr1^EC-KO^* and *Cnr1^EC-WT^* mice after 4 weeks of WD and performed RNA sequencing. Principal component analysis showed distinct clustering of *Cnr1^EC-KO^* and *Cnr1^EC−WT^* samples (Figure 3A). We found 303 differentially expressed genes (DEGs) between the two groups, of which 129 were downregulated and 174 were upregulated in *Cnr1^EC-KO^* compared to *Cnr1^EC-WT^* ECs. Notably, the expression of several pro-inflammatory cytokines and chemokines, such as *Il6*, *Ackr3*, *Cxcl12*, and *Ccl2*, was significantly lower in *Cnr1^EC-KO^* ECs, consistent with a less inflammatory phenotype (Figure 3B). Conversely, genes associated with the cellular matrix, including *Itga8* and *Itga3*, and the transcription factor *Prdm16*, which plays a critical role in maintaining endothelial function and supporting arterial flow recovery,^23^ were upregulated in *Cnr1^EC-KO^* ECs. Taken together, these findings suggest a complex network of transcriptomic changes that may underlie the CB1-dependent regulation of EC homeostatic function.

**Figure 3.**
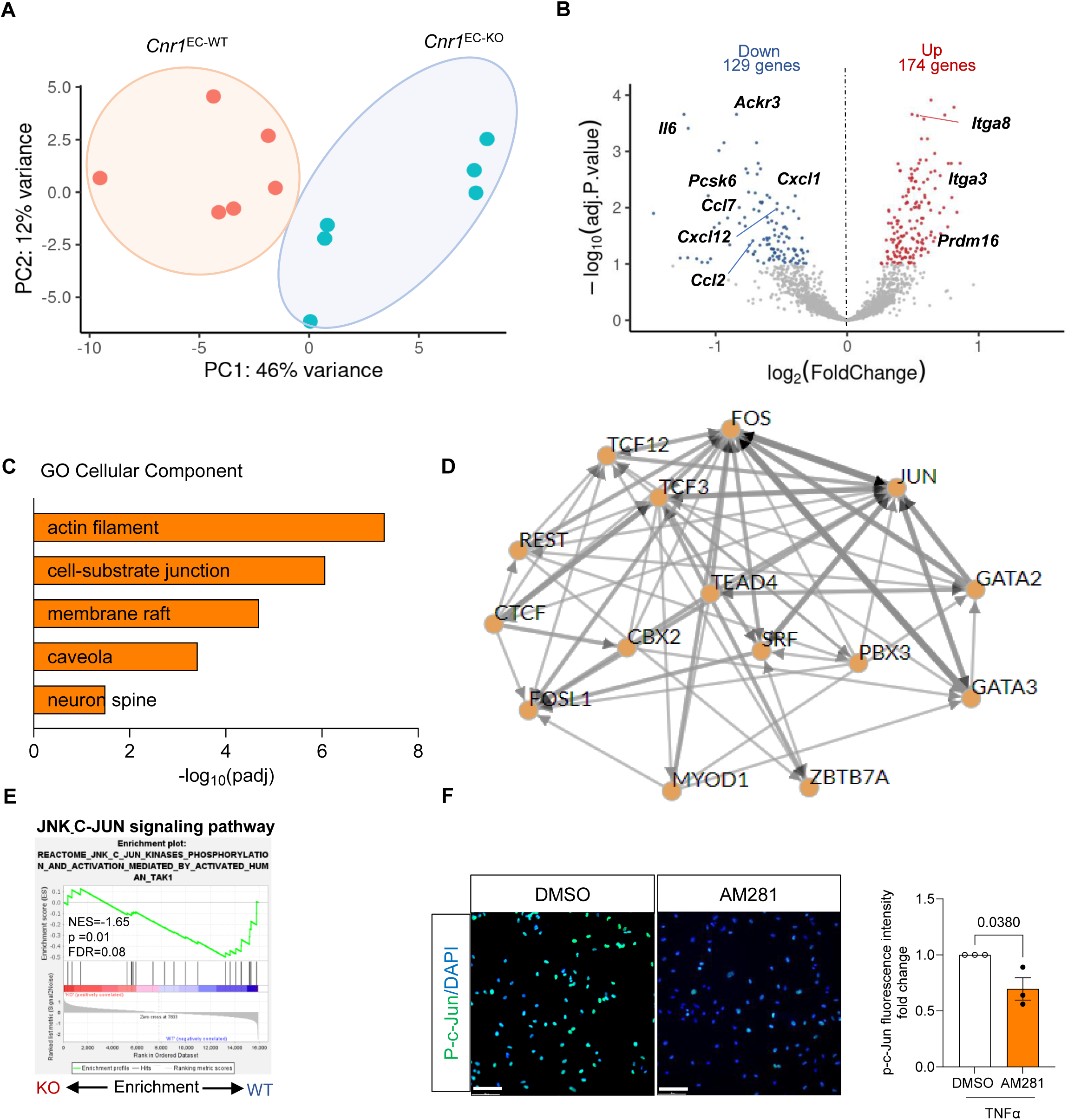
Transcriptomic profile of aortic endothelial *Cnr1* signalling. Aortic ECs (sorted as CD45^−^CD107a^low^CD31^high^) were isolated from female *Cnr1^EC-WT^* and *Cnr1^EC-KO^* mouse aortas (n=6) after 4 weeks of WD for RNA sequencing. (**A**) Principal component analysis (PCA) to visualize the heterogeneity of transcriptomic profiles between the two groups. (**B**) Volcano plot showing differentially expressed genes (DEGs) with 174 significantly up- and 129 down-regulated genes in female *Cnr1^EC-WT^* and *Cnr1^EC-KO^* mice. (**C**) Significantly regulated pathways in *Cnr1* deficient ECs, based on GO analysis. (**D**) CHEA3 was used to predict the top 15 transcription factor (TF) -co-regulatory networks modulated by endothelial *Cnr1* deficiency. (**E**) Pathway enrichment (GSEA) with normalized enrichment score (NES) and false discovery rate (FDR) q-value. NES below −1 or above 1 and an FDR q-value below 0.25 is considered significant. (**F**) Phospho-c-Jun immunostaining (green) of HAoECs pretreated with vehicle or 1 μM AM281 for 5 min prior to 30 min TNFα stimulation. Nuclei were stained with DAPI (blue); scale bar, 20 μm. Data are shown as mean ± s.e.m. and unpaired Student’s t-test was performed.

Further gene ontology (GO) analysis revealed that the transcriptomic signature regulated by endothelial *Cnr1* deficiency affects key cellular components, including membrane raft (GO:0045121; padj = 2.05e^−05^) and caveola (GO:0005901; padj = 0.0003), suggesting that CB1 may affect endothelial lipid raft and caveolae-dependent signaling (Figure 3C). To identify putative upstream transcription factors (TFs) of the transcriptomic signature in *Cnr1*-deficient ECs, ChIP-X Enrichment Analysis 3 (CHEA3) was performed (Figure 3D). *Fos* and *Jun* emerged as the primary TFs regulating the DEGs network. These TFs are known as activator protein 1 (AP-1) regulators and mediate various cellular processes such as cytokine and chemokine expression, as well as cell migration and differentiation. In addition, GSEA revealed enrichment of DEGs in the JNK_c-JUN, Il6_JAK_Stat3, inflammatory, and NFκB signaling pathways (Figure 3E, Supplemental Figure S4A). The regulation of c-JUN activation by CB1 was further investigated in HAoECs treated with the CB1 antagonist AM281, which significantly inhibited TNFα-induced nuclear translocation of phosphorylated c-JUN (Figure 3F and Supplemental Figure S4B).

To validate our findings, we investigated the regulation of key pro-inflammatory markers among the identified DEGs in HAoECs after transfection with *CNR1* or scrambled siRNA. A knockdown efficiency of approximately 80 % was achieved with *CNR1* siRNA (Supplemental Figure S4C). In *CNR1*-silenced HAoECs, the expression of the pro-inflammatory genes *CXCL8*, *CCL2* and *ICAM1* was significantly attenuated. (Supplemental Figure S4D), confirming a reduced pro-inflammatory phenotype in the absence of endothelial CB1 signaling. Because increased vascular inflammation disrupts vascular homeostasis and induces endothelial oxidative stress,^24^ we tested whether CB1 signaling affects ROS production. Flow cytometry analysis revealed a reduction in TNF-α-stimulated ROS production in HAoECs after *CNR1* knockdown (Supplemental Figure S4E). To further confirm that CB1 signaling promotes endothelial inflammation, we treated HAoECs with the synthetic CB1 agonist ACEA in LSS culture conditions and found that CB1 stimulation prevented the atheroprotective effects of LSS exposure. While LSS downregulated the expression of *ICAM1*, vascular cell adhesion molecule-1 (*VCAM1*), and glycolytic enzyme *PFKFB3* compared to the static condition, this effect was attenuated by ACEA (Supplemental Figure S4F). In addition, ACEA prevented LSS-induced upregulation of *KLF2* and *NOS3* (Supplemental Figure S4F). Comparable expression levels were detected in vehicle- and ACEA-treated ECs under static conditions, suggesting that CB1 activation under shear stress inhibited the LSS-mediated anti-inflammatory phenotype. To support these findings, an adhesion assay was performed with HAoECs perfused with labeled THP-1 monocytes in the presence or absence of the CB1 agonist ACEA under LSS. A significantly increased number of adherent monocytes was observed when stimulating HAoECs with ACEA compared to the vehicle control (Supplemental Figure S4G-H). Overall, our results demonstrated that impaired CB1 signaling exhibits an anti-inflammatory phenotype in both human and murine endothelial cells.

### Endothelial *Cnr1* deficiency reduces atherosclerotic plaque formation in female mice

To address whether the reduced inflammatory phenotype in the absence of endothelial CB1 translates into reduced atherosclerotic lesion development, we subjected *Cnr1^EC-KO^* and *Cnr1^EC-WT^* mice to a WD for 4 weeks or 16 weeks, to induce early and advanced stages of atherosclerosis, respectively. At the 4-week time point, *Cnr1^EC-KO^* mice exhibited plaque sizes comparable to those of *Cnr1^EC-WT^* mice in the aortic roots and descending aortas, whereas significantly smaller plaques were found in the aortic arch of female *Cnr1^EC-KO^* mice compared to *Cnr1^EC-WT^* mice (Figure 4A-F). After 16 weeks of atherogenic diet, female *Cnr1^EC-KO^*mice exhibited smaller atherosclerotic plaques in the aortic roots compared to corresponding *Cnr1^EC-WT^* controls (Figure 4G-I), suggesting that loss of endothelial *Cnr1* ameliorates atherosclerosis in female mice. To exclude any effects of the *loxP* insertion in *Cnr1^flox/flox^* mice on the atherosclerotic phenotype*, Apoe^−/−^Cnr1^flox/flox^* mice were used as an additional control group (Supplemental Figure S5A). No impact of endothelial *Cnr1* deficiency on lesion development was observed in male mice, possibly because lesions develop more slowly in male mice compared to female mice. In support of this hypothesis, the difference in females was only evident at the predilection sites of the aortic arches at 4 weeks WD or at an advanced stage at 16 weeks WD. Notably, measurement of plasma endocannabinoid levels in male and female *Apoe^−/−^* mice after 4 weeks of WD revealed comparable levels of 2-arachidonoylglycerol, anandamide, oleoylethanolamide, and palmitoylethanolamide, excluding that the sex-specific effect of endothelial *Cnr1* deficiency on atherosclerotic plaque development was generated by differences in circulating endocannabinoid levels (Supplemental Figure S5B).

**Figure 4.**
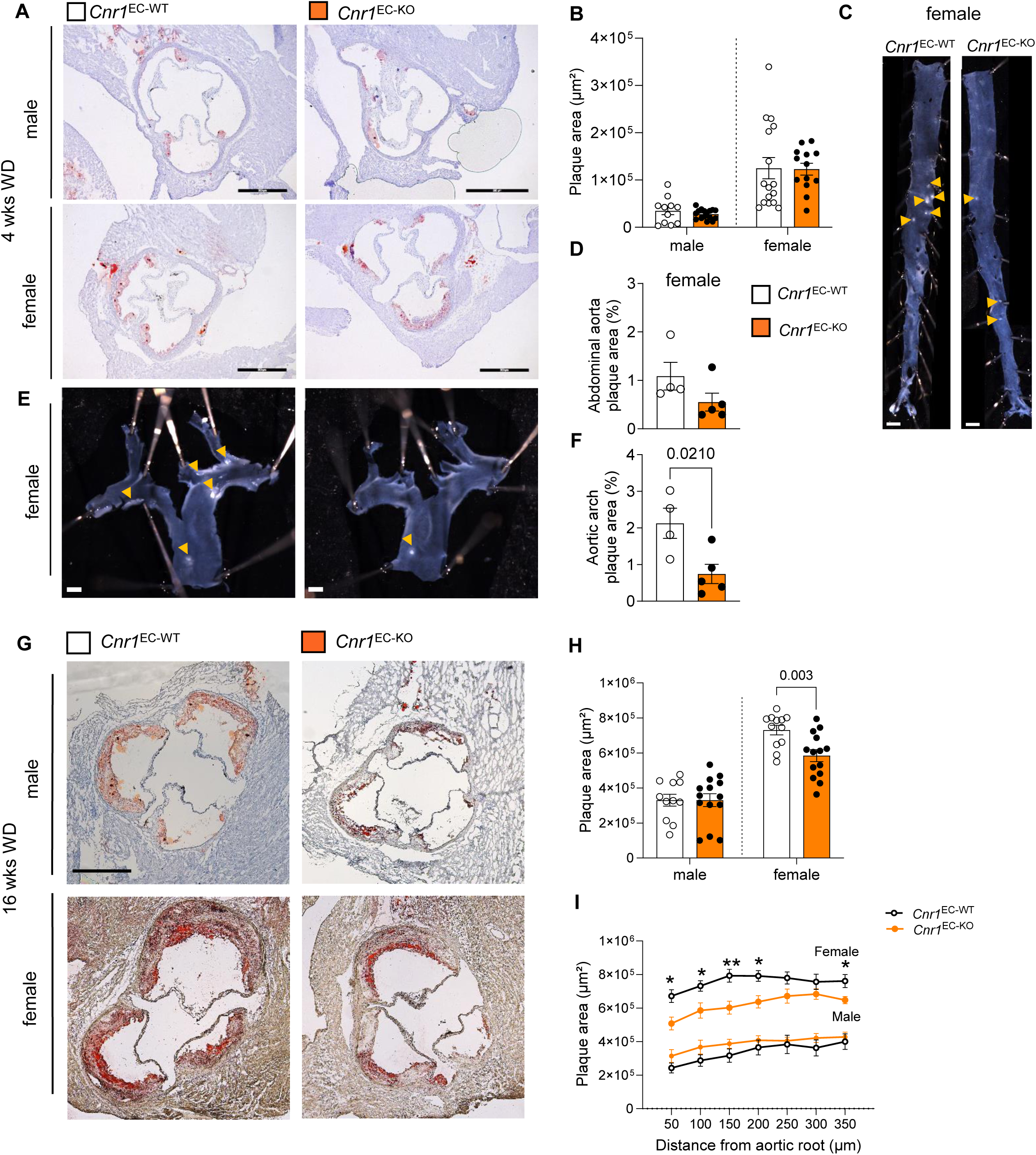
Effect of endothelial *Cnr1* deficiency on atherosclerotic plaque development. (A) Representative Oil-Red-O (ORO) stainings of aortic roots from male (n=11-16) and female (n=13-16) *Cnr1^EC-WT^* and *Cnr1^EC-KO^* mice after 4 weeks of WD; Scale bar, 500 μm. (**B**) Quantification of absolute lesion area. (**C-F**) Representative images and analysis of arch and descending aorta lesion area normalized to vessel area from female *Cnr1^EC-WT^* and *Cnr1^EC-KO^* mice (n = 4-5) after 4 weeks of WD. Scale bar, 1 mm. (**G**) Representative ORO staining of aortic roots of male (n=7-14) and female (n=8-14) *Cnr1^EC-WT^* and *Cnr1^EC-KO^* after 16 weeks of WD; Scale bar, 500 μm. (**H**) Quantification of absolute lesion area. (**I**) Plaque area per aortic root section. Data are shown as mean ± s.e.m. and exact p values are indicated or shown as **p<0.01 and *p<0.05. Unpaired Student’s t-test (**B**, **D**, **F**, **H**, **I**) was performed.

Surprisingly, male and female *Cnr1^EC-KO^* mice had significantly higher circulating plasma cholesterol levels compared to the *Cnr1^EC-WT^*control group (Supplemental Figure S5C), especially in the LDL fraction (Supplemental Figure S5D). Because hypercholesterolemia promotes arterial inflammation and monocyte recruitment,^1,25^ this finding was unexpected in light of the reduced plaque size observed in female mice. However, no significant differences in circulating leukocyte counts were observed between the *Cnr1^EC-KO^* and *Cnr1^EC-WT^* groups (Supplemental Figure S5E). We hypothesized that the increase in circulating plasma cholesterol may be a consequence of reduced tissue lipid uptake. Reduced aortic lipid uptake may explain a reduced plaque development despite elevated plasma cholesterol levels. To characterize advanced aortic root lesions after 16 weeks of WD, we assessed intracellular lipid droplets, necrotic core size, collagen and macrophage content. Significant differences were found in the plaque composition of female *Cnr1^EC-KO^* mice, in particular a significant reduction in relative lipid plaque content and an increase in collagen content (Supplemental Figure S6A-D). These findings suggest a shift towards a more stable plaque phenotype, possibly due to a reduced aortic lipid accumulation.

### Endothelial *Cnr1* deficiency improves metabolic parameters

We found that the depletion of *Cnr1* in ECs was sufficient to attenuate the body weight gain over 16 weeks WD in male and female mice (Figures 5A and Supplemental Figure S7A), as previously reported in global *Cnr1* deficient mice subjected to a high fat diet. *Cnr1^EC-KO^* mice had less white and brown fat mass (Figures 5B and Supplemental Figure S7B). Histological analysis of female epididymal white adipose tissue at 4 weeks WD revealed a significant reduction in adipocyte size of *Cnr1^EC-KO^*mice, whereas the relative lipid droplet content in brown adipose tissue (BAT) was not different (Figures 5C-D). At the same time point, we detected higher mRNA expression of *Gpihbp1*, a capillary endothelial cell protein that facilitates lipolysis of triglyceride-rich lipoproteins, as well as other lipolysis-related genes in the BAT of *Cnr1^EC-KO^*male and female mice (Figure 5E and Supplemental Figure S7C). Increased capillary endothelial GPIHBP1 expression in the BAT of female *Cnr1^EC-KO^* mice was confirmed by immunostaining (Figure 5F). The transcription factor *Prdm16*, which is involved in adipose tissue browning and thermogenesis, was also upregulated in the BAT of *Cnr1^EC-KO^* mice (Figure 5E and Supplemental Figure S7C). This prompted us to further investigate the underlying EC-specific transcriptional changes. To this end, we sorted ECs from the BAT of female C*nr1^EC-KO^* and C*nr1^EC-WT^* mice at 4 weeks WD for RNA sequencing. The GO analysis revealed that the pathways regulated by endothelial CB1 in the BAT were related to mitochondrial respiration and ATP biosynthesis, suggesting increased mitochondrial activity in the BAT of C*nr1^EC-KO^* mice (Figures 5G). Examination of the liver phenotype in female C*nr1^EC-KO^* mice revealed reduced hepatic accumulation of lipid droplets (Figures 5H) and decreased hepatic cholesterol levels at 4 weeks WD (Figures 5I), while plasma bile acid levels were not affected by endothelial *Cnr1* deficiency when analyzed in both males and females (Supplemental Figure S7D). Liver qPCR analysis at the same time point showed a significant increase in the expression of the fatty acid transporter *Cd36* and the marker *Acadm* (acyl-CoA dehydrogenease medium chain) linked to hepatic β-oxidation in female C*nr1^EC-KO^* mice (Figures 5J). In addition, an intraperitoneal glucose tolerance test (ipGTT) revealed improved glucose disposal rates in C*nr1^EC-KO^*male and female mice compared to the age-matched C*nr1^EC-WT^* mice (Supplemental Figure S7E-F).

**Figure 5.**
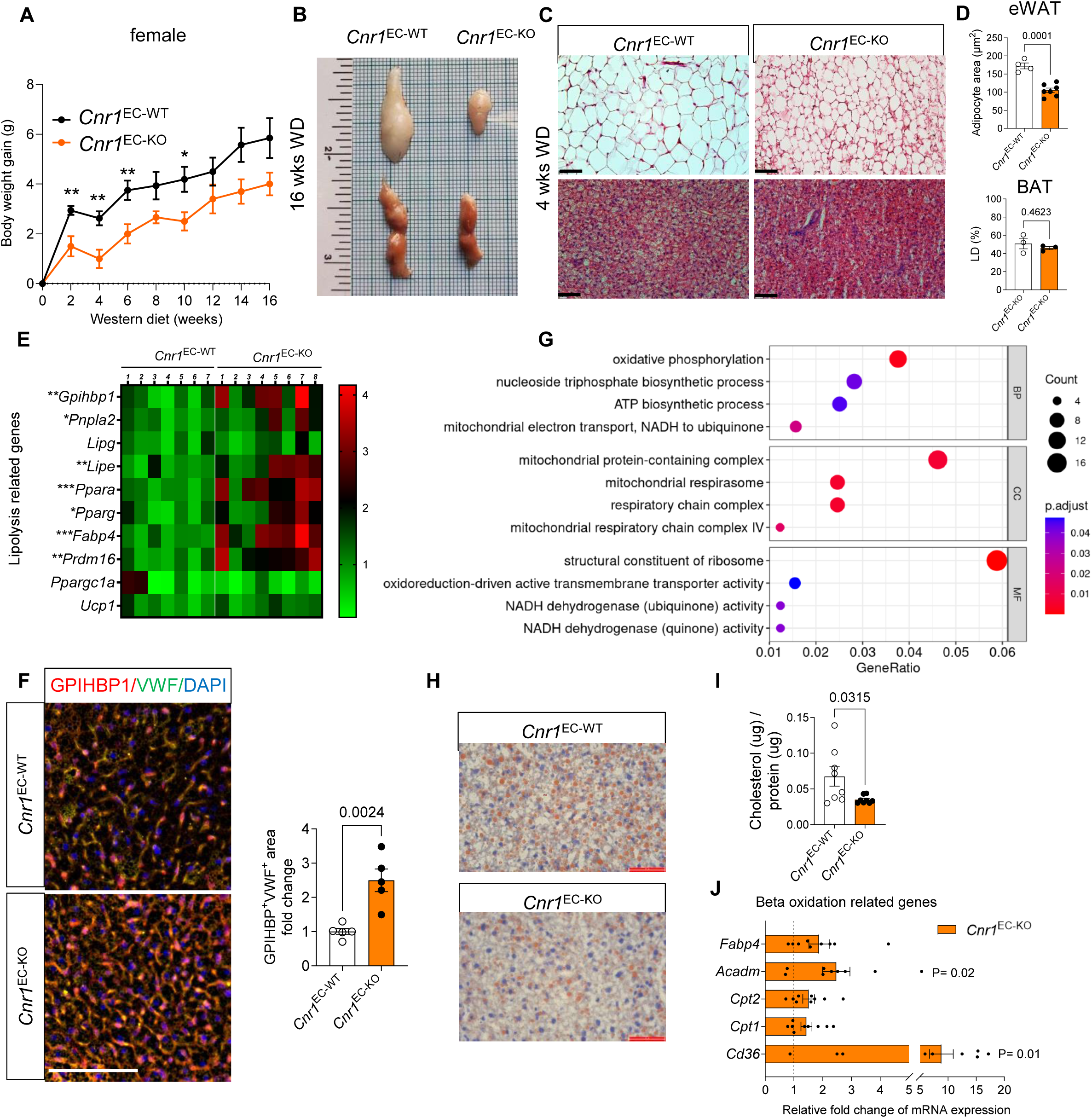
Impact of endothelial *Cnr1* deficiency on lipid metabolism. (**A**) Body weight gain over 16 weeks WD in female *Cnr1^EC-WT^* and *Cnr1^EC-KO^* mice (n=7-8). (**B**) Representative images of epididymal white (eWAT, upper) and brown adipose tissue (BAT, lower) from female mice after 16 weeks WD. (**C**) H&E staining of eWAT and BAT from female *Cnr1^EC-WT^* and *Cnr1^EC-KO^* after 4 weeks of WD. Scale bar, 500 μm, and (**D**) corresponding quantification of adipocyte size in eWAT and relative lipid droplet (LD) content in BAT. (**E**) Gene expression analysis (RT-qPCR) in BAT of female *Cnr1^EC-WT^* and *Cnr1^EC-KO^* mice after 4 weeks WD (n = 7-8). (**F**) Co-staining of GPIHBP1 (red) and von Willebrand factor (vWF, green; double positive, yellow) for capillary vessels in BAT of female *Cnr1^EC-WT^* and *Cnr1^EC-KO^* mice after 4 weeks of WD. Scale bar, 100 μm. Quantification of GPIHBP1^+^VWF^+^ area and calculated as fold change to WT. (**G**) RNA sequencing and GO analysis of regulated pathways in sorted BAT ECs (CD45^−^CD31^+^) from female *Cnr1^EC-KO^* mice after 4 weeks WD (n=7). (**H**) Representative liver ORO staining and (**I**) liver cholesterol levels in female *Cnr1^EC-WT^* and *Cnr1^EC-KO^* mice after 4 weeks WD (n=8). Scale bar, 50 μm. (**J**) Gene expression analysis (RT-qPCR) in livers of female *Cnr1^EC-KO^* and *Cnr1^EC-WT^* mice after 4 weeks WD (n = 8-9). Data are shown as mean ± s.e.m. and two-tailed Student’s t-test was performed. ***p<0.001; **p<0.01; *p<0.05

### Endothelial *Cnr1* deficiency reduces caveolae-mediated LDL uptake

To confirm our hypothesis that the increase in circulating plasma cholesterol and reduced plaque lipid content in *Cnr1*^EC-KO^ mice may be a consequence of reduced tissue lipid uptake, we perfused carotid arteries from female C*nr1^EC-KO^* and C*nr1^EC-WT^* mice isolated at 4 weeks WD with fluorescently labeled native LDL (Dil-LDL).^26^ Two-photon laser scanning microscopy (TPLSM) revealed that lack of *Cnr1* in endothelial cells resulted in significantly reduced retention of LDL particles across the endothelial layer (Figure 6A-B). To further investigate the underlying CB1-dependent regulation of endothelial LDL uptake, we screened for candidate receptor expression levels in our RNA sequencing data of sorted C*nr1^EC-KO^* and C*nr1^EC-WT^* aortic endothelial cells and found a significantly lower expression level of *Acvrl1*, which encodes the protein activing A receptor like type 1, also termed activing-like kinase receptor (ALK1, Figure 6C). We did not find any transcriptional regulation of other endothelial LDL receptors including the canonical LDLR (Supplemental Figure S8A). In agreement with the transcriptomic data, no differences in the surface expression levels of LOX-1, SRB1, SRA1, and CD36 were found between C*nr1^EC-KO^* and C*nr1^EC-WT^* aortic endothelial cells (Supplemental Figure S8B-C).

**Figure 6.**
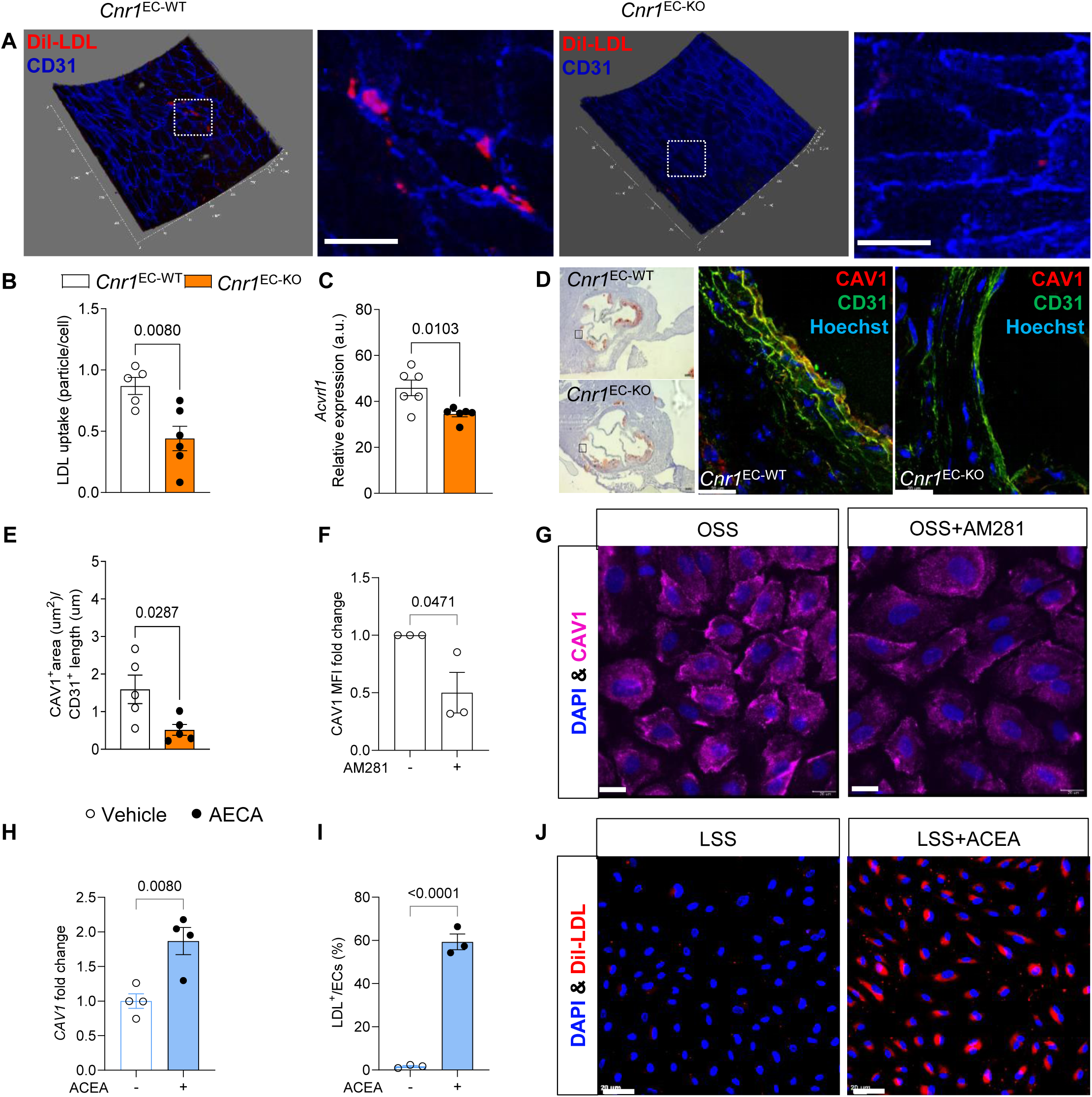
Role of CB1 in endothelial LDL transport. (**A**) Representative TPLSM 3D images of Dil-LDL particles in ex vivo perfused carotid arteries from female *Cnr1^EC-WT^* and *Cnr1^EC-KO^* mice after 4 weeks WD; CD31 was used to stain ECs. Scale bar, 20 μm. (**B**) Quantification of Dil-LDL normalized to EC number (n=5-6 mice per group). (**C**) Endothelial ALK1 encoding gene expression retrieved from the RNA sequencing data of sorted aortic *Cnr1^EC-WT^* and *Cnr1^EC-KO^* ECs (n=6). (**D**) Representative confocal images of caveolin-1 (CAV1, red) and CD31 (green) staining with bright field overview images of consecutive aortic root sections of *Cnr1^EC-WT^* and *Cnr1^EC-KO^* mice after 4 weeks WD (scale bar, 20 μm) and (**E**) quantification of endothelial CAV1 staining normalized to EC length (n=5 mice). (**F, G**) Quantification of CAV1 mean fluorescence intensity (MFI) and representative images of HAoECs treated with 1 μM AM281 or vehicle under OSS for 24 h followed by 1ug/ml Dil-LDL treatment for 30 min. Scale bar, 20 μm. (**H**) *CAV1* mRNA expression (RT-qPCR) in HAoECs treated with 1 μM ACEA or vehicle under LSS for 24 h. (**I**, **J**) Quantification of Dil-LDL uptake (calculated as percentage of all cells) and representative images of HAoECs preincubated with 1 µM ACEA or vehicle under LSS for 24 h, followed by 1 µg/mL Dil LDL treatment for 90 min. Scale bar, 50 μm. Unpaired Student’s t-test (**B,C,E,F,H,I**) was performed.

In support of CB1-dependent regulation of LDL transcytosis, the GO analysis of the aortic EC transcriptomic data indicated a regulation of membrane rafts and caveola in *Cnr1* deficient endothelial cells (Figure 3C). CAV1, a major component of caveolae, co-localizes with ALK1 within caveolae-enriched domains of ECs.^5^ Genetic deficiency of *Cav1* has been shown to inhibit aortic DiI-LDL uptake in isolated aortas and atherosclerosis despite elevated total cholesterol levels.^27^ We performed co-immunostaining of endothelial cells and CAV1 in aortic root sections from C*nr1^EC-KO^* and C*nr1^EC-WT^* mice at the 4 week WD time point. CAV1 expression in C*nr1^EC-KO^* endothelial cells was significantly reduced in female mice (Figure 6D-E), providing a possible mechanism for reduced endothelial LDL uptake in C*nr1^EC-KO^* mice. Notably, no difference was observed in male mice (Supplemental Figure S8D).

To further investigate the regulation of CAV1 and LDL uptake by CB1, HAoECs were treated with the CB1 antagonist AM281 under OSS, which resulted in significantly reduced CAV1 expression (Figure 6F-G). Conversely, when HAoECs were subjected to LSS, activation of CB1 with the agonist ACEA resulted in increased mRNA expression of *CAV1* and increased DiI-LDL uptake (Figure 6H-J). Taken together, these results support a direct regulation of endothelial caveolae-mediated LDL uptake by CB1 through modulation of *CAV1* gene expression.

### Endothelial CB1 induces caveolae-mediated LDL uptake via cAMP-PKA signaling

CB1 has been described in various cell lines as a G_i_ protein-coupled receptor that decreases intracellular cyclic adenosine monophosphate (cAMP) upon agonist binding.^28–30^ We first validated the CB1-dependent regulation of intracellular cAMP levels in response to CB1 activation or antagonism, respectively, in a HEK293 reporter cell line stably transfected with a cAMP-luciferase reporter plasmid and the human *CNR1* cDNA. The CB1 antagonist AM281 induced a significant increase in intracellular cAMP levels, whereas the CB1 agonist ACEA decreased intracellular cAMP levels, which was most evident when the adenylate cyclase activator forskolin was added shortly after agonist or antagonist administration (Supplemental Figure S8E). In HAoECs, endogenous cAMP levels were measured by ELISA, which showed that AM281 dose-dependently increased intracellular cAMP levels under static conditions, while ACEA showed only modestly decreased endogenous cAMP levels, possibly due to intrinsic activation of CB1 by endocannabinoids under basal conditions (Figure 7A). These data confirm that pharmacological inhibition of endothelial CB1 signaling leads to an increase in intracellular levels of cAMP.

**Figure 7.**
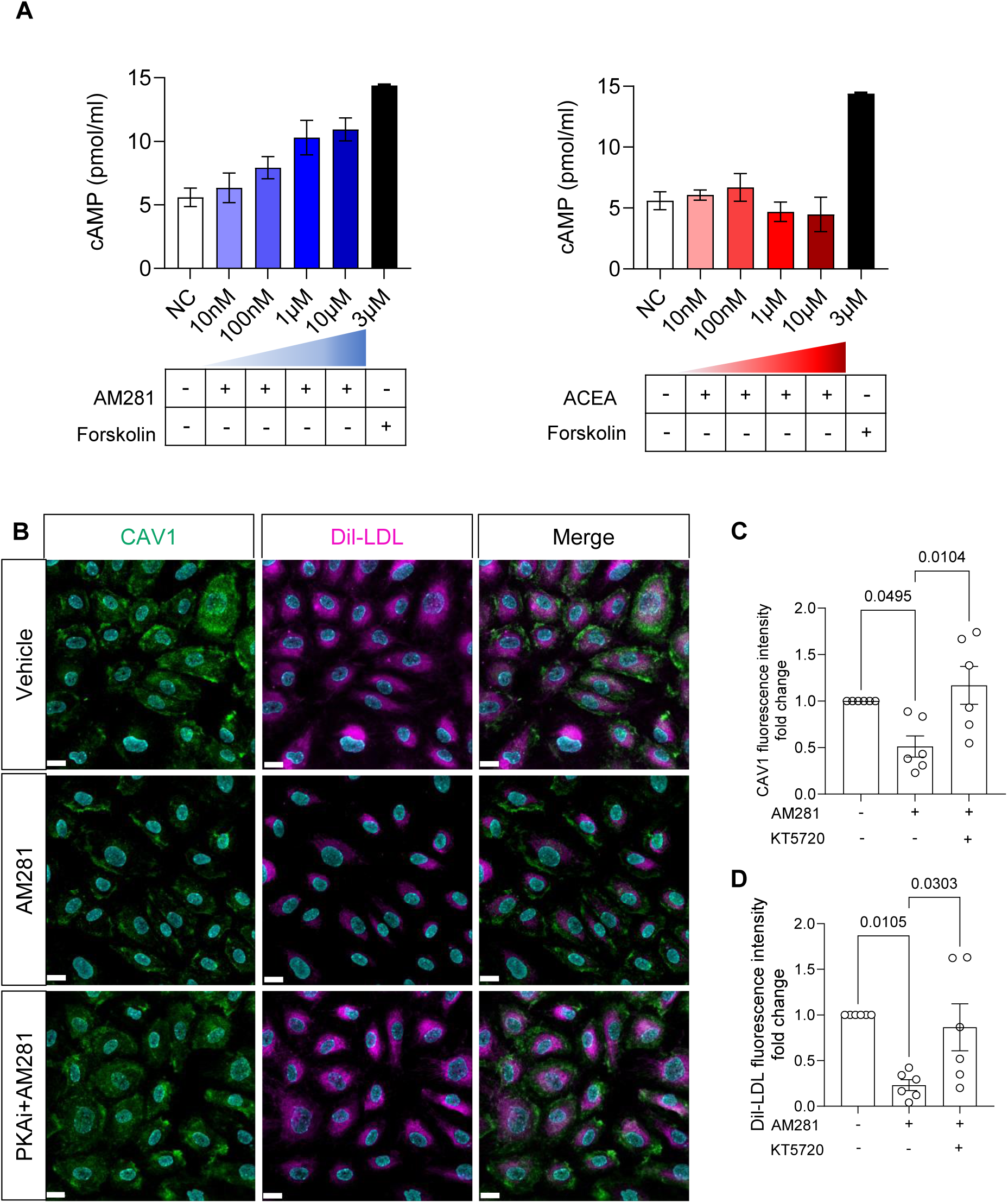
Role of PKA in CB1-dependent regulation of endothelial CAV1 expression and LDL transport. (**A**) cAMP levels in HAoECs 20 min after treatment with AM281, ACEA, or adenylate cyclase activator forskolin, assessed via ELISA (n=2). (**B**) Representative immunofluorescence analysis of CAV1 expression and Dil-LDL uptake in HAoECs treated with 1 μM AM281 alone or in the presence of 1 μM PKA inhibitor (KT5720) or vehicle (DMSO) under OSS for 24 h. Scale bar, 20 μm. Quantification of (**C**) CAV1 fluorescence intensity and (**D**) LDL uptake fold change (n=6 independent experiments). Data are shown as mean ± s.e.m. and one-way ANOVA with Tukey correction was performed.

It has been previously reported that the activation of protein kinase A (PKA) leads to a reduction in CAV1 expression in Chinese hamster ovary cells.^31^ PKA is activated upon binding of cAMP to its regulatory subunits.^32^ To clarify whether endothelial CAV1 expression is regulated by CB1 via cAMP-PKA signaling, HAoECs cells were treated with KT5720, a PKA inhibitor, before the addition of AM281 under OSS conditions. The reduction in endothelial CAV1 expression by AM281 treatment was prevented by the addition of KT5720 (Figure 7B-C). Similarly, the reduction in LDL uptake by AM281 treatment was prevented by pretreatment with KT5720 (Figure 7D), suggesting that the reduction in CAV1 requires PKA activation in ECs. Notably, all the above-described *in vitro* experiments were performed with primary human cells from female donors. When the same experiments were repeated with HAoECs from an age-matched male donor, no significant effects on CAV1 expression were observed, only reduced LDL uptake (Supplemental Figure S9A-B).

### Chronic peripheral CB1 antagonism reduces atheroprogression, endothelial CAV1 and ICAM1 expression in female Ldlr^−/−^ mice

To clarify whether the peripheral CB1 antagonist JD5037 would reproduce an atheroprotective phenotype as observed with endothelial *Cnr1* deficiency, we first subjected *Ldlr^−/−^*mice to a Western diet for 8 weeks to induce atherosclerotic lesions, followed by an additional 8-weeks of JD5037 treatment in parallel with continuous WD (Figure 8A). We used the *Ldlr^−/^*^−^ atherosclerotic mouse model for this therapeutic approach because it more closely mirrors the lipid profile in humans compared to the *Apoe^−/−^* model.^33^ Consistent with our previous study focusing on atheroprotective effects of peripheral CB1 antagonism on myeloid cells at an earlier time point, chronic administration of the peripheral CB1 antagonist JD5037 resulted in significantly less body weight gain,^16^ but plasma total cholesterol and triglyceride levels were not different after 16 weeks of WD (Supplemental Figure S10A-D).

**Figure 8.**
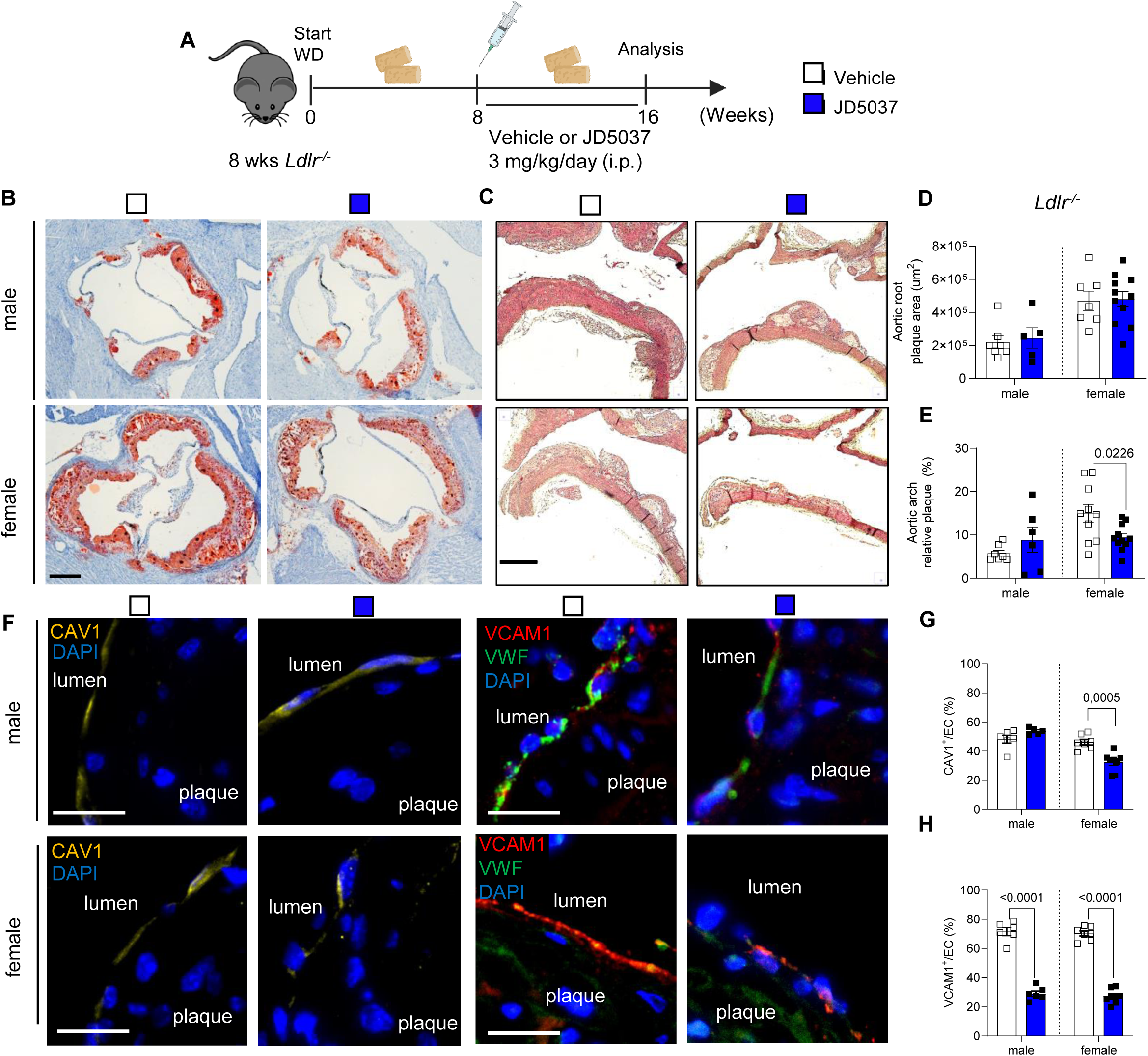
Effect of chronic peripheral CB1 antagonist administration on atherosclerosis and CAV1 expression. (**A**) Schematic experimental design. *Ldlr^−/−^* mice were fed with WD for 16 weeks and received daily intraperitoneal injections of JD5037 (3 mg/kg) or vehicle for the last 8 weeks. (**B**) Representative ORO-stained images of aortic root cross-sections (scale bar, 500 μm) and (**C**) H&E-stained aortic arch longitudinal sections (scale bar, 500 μm). (**D**) Quantification of absolute lesion areas in aortic roots (n=5-9) and (**E**) aortic arches (n=7-10). (**F**) Representative images of CAV1 (yellow), VCAM (red) and vWF (green) in aortic arch plaques; nuclei were stained with DAPI (scale bar, 20 μm). (**G-H**) Quantification of CAV1 and VCAM1 positive ECs. Each data point represents a mouse, and all data are expressed as mean ± s.e.m. P values were obtained using an unpaired Student’s t-test. Males and females were analysed independently.

No significant effects of JD5037 treatment on plaque size progression were observed in the aortic root compared to the corresponding sex-matched vehicle group. However, a reduction in plaque progression in the aortic arch was observed in female *Ldlr^−/^*^−^ mice treated with JD5037; this effect was not seen in male mice (Figure 8B-E). Consistent with the observations in C*nr1^EC-KO^* mice, reduced plaque size in female JD5037-treated mice was accompanied by decreased levels of CAV1, ICAM1, and VCAM1 in aortic endothelial cells (Figure 8F-H and Supplemental Figure S10E). A significant reduction in endothelial VCAM1 expression was also observed in male JD5037-treated mice compared to the vehicle group (Figure 8H). These results suggest that peripheral CB1 antagonism blocks the proatherogenic effects of CB1 in arterial endothelial cells, with more pronounced effects in female mice.

## DISCUSSION

In this study, we provide evidence that CB1 is expressed by human plaque endothelial cells and is upregulated by atherogenic shear stress. We demonstrated that the loss of endothelial CB1 affects vascular tone and reverses endothelial dysfunction after atherogenic diet feeding. Notably, we found that the absence of endothelial CB1 reduces arterial LDL infiltration and improves metabolic parameters in brown and white adipose tissue and liver. In addition, the peripheral CB1 antagonist JD5037 provided therapeutic benefit by limiting plaque progression and endothelial inflammation. These findings reveal a pivotal role for endothelial CB1 as a central regulator linking biomechanical and inflammatory pathways with LDL translocation across the endothelial layer. Thus, antagonizing peripheral CB1 signaling may reduce cholesterol uptake in the arterial wall, thereby contributing to pleiotropic beneficial effects of CB1 blocking in atherosclerosis and related metabolic alterations.

We determined the expression of CB1 in human and mouse atherosclerotic vessels at the mRNA level due to the lack of specific antibodies to detect CB1 at the protein level.^34^ In human plaque single cell RNA sequencing data from the Munich Vascular Biobank,^18^ we detected *CNR1* only in ECs, probably because the sensitivity of analysis was insufficient to detect *CNR1* expressed by other plaque cell types. Remarkably, we found that endothelial CB1 expression is regulated by shear stress, revealing an increased receptor expression in atheroprone regions of mouse aortas and in HAoECs subjected to OSS. Whether endothelial CB1 itself may serve as a mechanosensitive receptor on the cell surface, responding to changes in shear stress patterns, remains unknown. It is also conceivable that *CNR1* is a downstream target of mechanosensitive transcription factors, which requires further investigation. Interestingly, the GO analysis of the transcriptomic profiling of murine aortic endothelial cells indicates that CB1 signaling is associated with the regulation of endothelial membrane raft and actin filament cellular components, which are involved in mechanosensing.^3^ It is possible that endothelial mechanosensing is affected in absence of endothelial CB1 due to a transcriptional regulation of mechanosensors. This may explain the observed differences in endothelial cell morphology in C*nr1^EC-KO^* aortas. Because shear stress response and changes in EC alignment depend on both the extracellular fibronectin matrix and the intracellular cytoskeletal F-actin,^35^ it is conceivable that endothelial CB1 affects endothelial morphology in response to blood flow through regulation of actin filament signaling. Furthermore, dynamic changes of actin filaments are closely linked to endothelial junctional VE-cadherin remodeling.^36^ Therefore, we can speculate that the changes in aortic permeability in the absence of endothelial CB1 may be due to alterations in actin filament signaling. The exact molecular mechanisms underlying this regulation remain to be elucidated.

CB1 signaling has been previously linked to the regulation of vascular tone, in particular in hypertensive conditions.^37^ Here, we found that the absence of endothelial CB1 increases flow velocity in atheroprone sites of the aorta, thereby sharing a similar phenotype as previously reported in *Cav1* deficient mice.^27^ This resulted in a preserved ejection fraction and fractional shortening after atherogenic diet feeding. Taken together, these findings support a central role for endothelial CB1 in the regulation of the homeostatic functions of the vasculature.

In atherogenic conditions, endothelial *Cnr1* deficiency resulted in a transcriptional downregulation of several proinflammatory markers, including *Il6*, *Ccl2*, *Cxcr12*, and *Ackr3*. The proinflammatory role of endothelial CB1 signaling was functionally confirmed by endothelial monocyte adhesion assays under flow conditions. Deficiency of ACKR3 in arterial ECs has been shown to reduce atherosclerotic lesions by limiting arterial leukocyte recruitment, which was linked to decreased NF-κB activity.^38^ GSEA and prediction of the top transcription factors modulated by endothelial C*nr1* deficiency revealed a regulation of the JNK/c-JUN and NF-κB pathways. JNK and NF-κB are key regulators of flow-dependent inflammatory gene expression in endothelial cells in atherosclerosis.^39–41^ These findings support that endothelial CB1 serves as an upstream regulator of JNK and NF-κB-dependent inflammatory gene expression, which is supported by previously published *in vitro* data.^42,43^ Furthermore, we provide evidence that endothelial CB1 regulates caveolae-mediated LDL uptake, which is supported by our GSEA findings linking CB1 to regulation of membrane raft and caveolae. While classical lipid receptors such as LDLR, SRB1, and LOX-1 were unaffected by endothelial C*nr1* deficiency, CAV1 and ALK-1, which are associated with caveolae-mediated LDL transcytosis pathways that are independent of the LDLR, were downregulated in C*nr1^EC-KO^* mice. Depletion of endothelial CAV1 or ALK-1 leads to impaired arterial LDL uptake,^6,8^ which is in agreement with the reduced plaque lipid content and functional LDL uptake in perfused carotids of *Cnr1*^EC-KO^ mice. The changes in endothelial CAV1 expression and LDL uptake by CB1 agonism or antagonism under flow culture conditions support a direct CB1-dependent regulation.

Activation of CB1, a G protein-coupled receptor, has been reported to cause G_i_ protein-dependent inhibition of adenylate cyclase, thereby reducing intracellular cyclic AMP production. In smooth muscle cells, stimulation of adenylate cyclase with forskolin has been reported to decrease *Cav1* mRNA expression.^44^ In addition, activation of PKA, which occurs when cAMP binds to its regulatory subunits,^32^ reduced CAV1 expression in Chinese hamster ovary cells.^31^ In agreement with a CB1-cAMP-PKA regulatory pathway of CAV1 expression in endothelial cells, we show that antagonism of CB1 increases cAMP levels and decreases CAV1 in HAoECs, whereas inhibiting PKA restored the CAV1 expression (Supplemental Figure S11). Notably, c-JUN/JNK appears among the top transcription factors with binding sites in the *CAV1* gene promoter according to genecards.org.

Remarkably, endothelial *Cnr1* deficiency resulted in a significant improvement of metabolic parameters in mice subjected to WD. The reduced adipose tissue mass and weight gain in *Cnr1^EC-KO^* mice was associated with an upregulation of *Gpihbp1* expression in WAT and BAT, indicating an increased lipoprotein lipase-mediated capillary lipid uptake and lipolysis. Moreover, *Cnr1*^EC-KO^ mice showed upregulated *Prdm16* expression, which is an important transcriptional regulator mediating brown fat differentiation. Other factors may also contribute to the striking metabolic phenotype observed in *Cnr1^EC-KO^* mice, including reduced lipid accumulation, improved glucose metabolism, and hepatic β-oxidation. The precise mechanisms involved in endothelial CB1 dependent regulation of metabolic processes in WAT, BAT and liver deserve to be investigated in more detail in future studies.

Of note, despite an improved adipose tissue and liver metabolism, *Cnr1^EC-KO^* mice had more elevated plasma cholesterol levels, as previously reported in *Cav1*-deficient mice.^45^ A common phenotype of endothelial *Cav1* and *Cnr1* deficiency is the reduced aortic LDL infiltration and reduced lesion progression.^6^ The systemic effects on plasma cholesterol levels were only observed in the *Apoe^−/−^*background, but not in *Ldlr^−/−^* mice when treating mice chronically with the peripheral CB1 antagonist, indicating a potential clinical benefit without undesired side effects on plasma cholesterol levels.

In the present study, the effects of endothelial *Cnr1* deficiency on atherosclerotic plaque size and endothelial CAV1 expression were only observed in female mice compared to males, whereas the metabolic effects were sex-independent. Remarkably, our own recent findings revealed a sex-specific effect of myeloid *Cnr1* deficiency, with atheroprotective effects and reduced macrophage proliferation only observed in male mice.^16^ Furthermore, treatment of *Ldlr^−/−^* mice with peripheral antagonist during early atherogenesis conferred atheroprotection only in male mice, whereas no difference was seen in females, which is opposite to the here observed protective effects of peripheral CB1 antagonism in females at advanced stage of plaque development. This indicates that sex differences in CB1 signaling are cell type- and stage-specific.

A limitation of this study is the unavailability of a CB1-specific antibody for detection at the protein level, which restricts the precise localization of the CB1 receptor within the endothelial cell membrane and potential interaction partners. Nevertheless, we have provided unprecedented insights into key regulatory functions of CB1 signaling in endothelial cells. Genetic deficiency or pharmacological inhibition of endothelial CB1 signaling conferred an atheroprotective phenotype in female mice with improved metabolic function in males and females, reduced vascular inflammation and diminished LDL entry into the artery wall. Results from human aortic endothelial cells reinforced the anti-inflammatory signaling associated with CB1 receptor silencing or antagonist treatment, while CB1 activation induced a proinflammatory phenotype, monocyte adhesion and EC LDL uptake. In conclusion, peripheral CB1 antagonists may hold promise as an effective therapeutic strategy for treating atherosclerosis and related metabolic disorders.

## Acknowledgements

We are grateful to the entire ZVH animal facility team for their continuous support and to Beat Lutz for providing *Cnr1^flox/flox^* mice. The graphical summary was created with BioRender.com.

## Author contributions

B.C. study design, experiments, data analysis, manuscript writing; A.P. *in vitro* shear stress experiments and immunostainings; G.L. transcriptomics data analysis; A.K. echocardiography; Y. W. and G.S. contributed to in vivo experiments; L.N. *en face in situ* hybridization; R.M. multi-photon imaging; Y.J. mouse aorta preparations and sectioning; S.R. and M.G. histology and *in vitro* experiments; A.F. *in vitro* signaling experiments; M.H. flow cytometric cell sorting; D.R. prime-Seq; Y.D. and M.v.d.S. endocannabinoid measurement; Z.L, N.S., L.M. human plaque single cell transcriptomic data; V.P. flow chamber adhesion assays; S.H. HPLC analysis; C.W. critical infrastructure and funding; S. H. study design, supervision and funding; R.G.P. study design, in vivo experiments, data analysis, supervision, manuscript editing; S.S. study design, supervision, funding, manuscript writing. All authors have read and commented on the manuscript and approved submission.

---

**Supplemental Figure S1.**
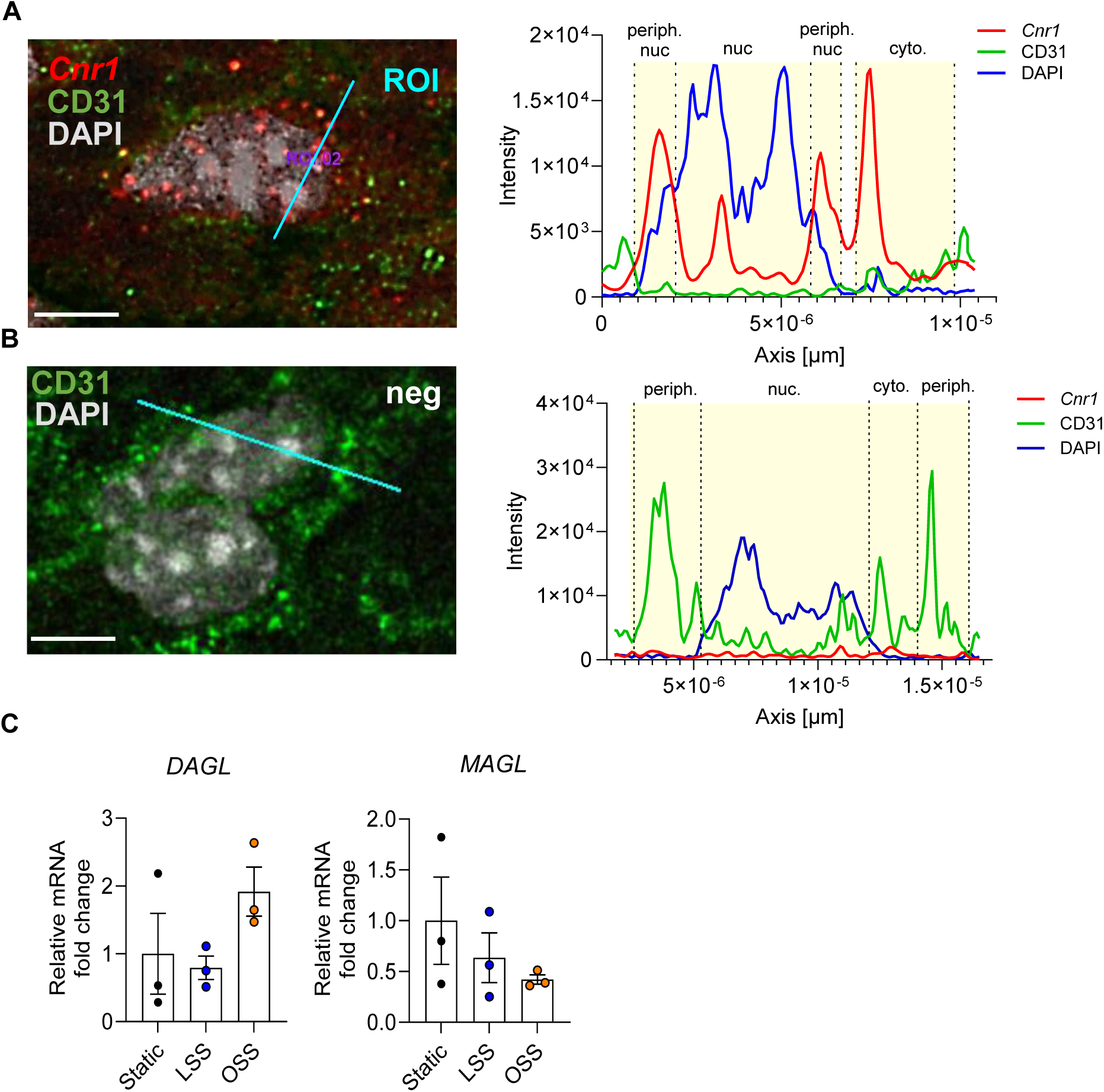
Intracellular localization of *Cnr1* in situ hybridization signal and endothelial regulation of endocannabioid-related genes by shear stress. (**A-B**) En face *in situ* hybridization was performed for *Cnr1* and corresponding negative control in thoracic aortas of 8-week-old *Apoe^−/−^* mice. The intensity of *Cnr1* (red), CD31 (green) and DAPI (grey) within endothelial cells in the region of interest (ROI) was quantified using IMARS, as presented on the right (scale bar, 5 μm). (**C**) Endocannabinoid synthesis and degradation enzyme *DAGL* and *MAGL* mRNA levels determined by RT-qPCR. The data are displayed as mean ± s.e.m., and each dot on the graph represents an independent experiment (n=3). One-way ANOVA followed by a post-hoc Tukey multi-comparison test was performed.

**Supplemental Figure S2.**
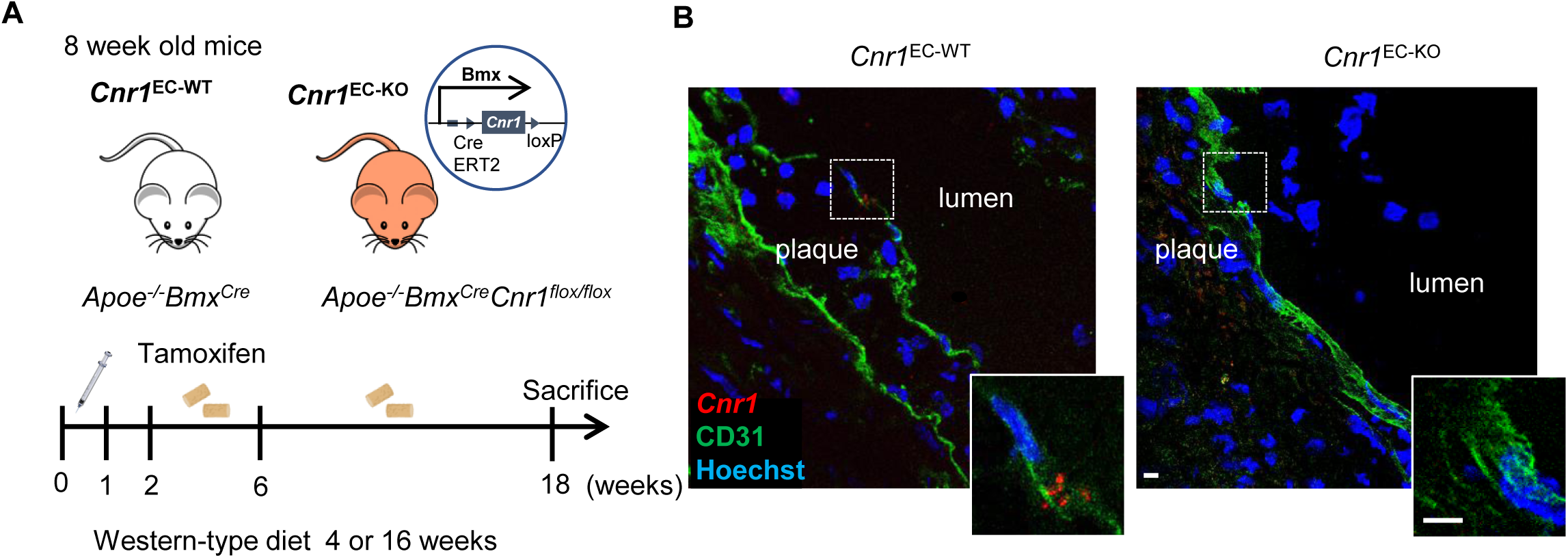
Validation of mouse model with selective endothelial *Cnr1* depletion. (**A**) Experimental scheme: 8 week old *Cnr1^EC-WT^* and *Cnr1^EC-KO^* mice received daily intraperitoneal injections of tamoxifen for 5 days and were subsequently fed with Western diet (WD) for a duration of 4 or 16 weeks. (**B**) Representative images of *in situ* hybridization for *Cnr1* (red) combined with CD31 (green) immunostaining for ECs and Hoechst 33342 staining (blue) for nuclei in aortic root sections of *Cnr1^EC-WT^* and *Cnr1^EC-KO^* mice after 4 weeks WD. Scale bar: 20 μm (overview) and 5 μm (insert).

**Supplemental Figure S3.**
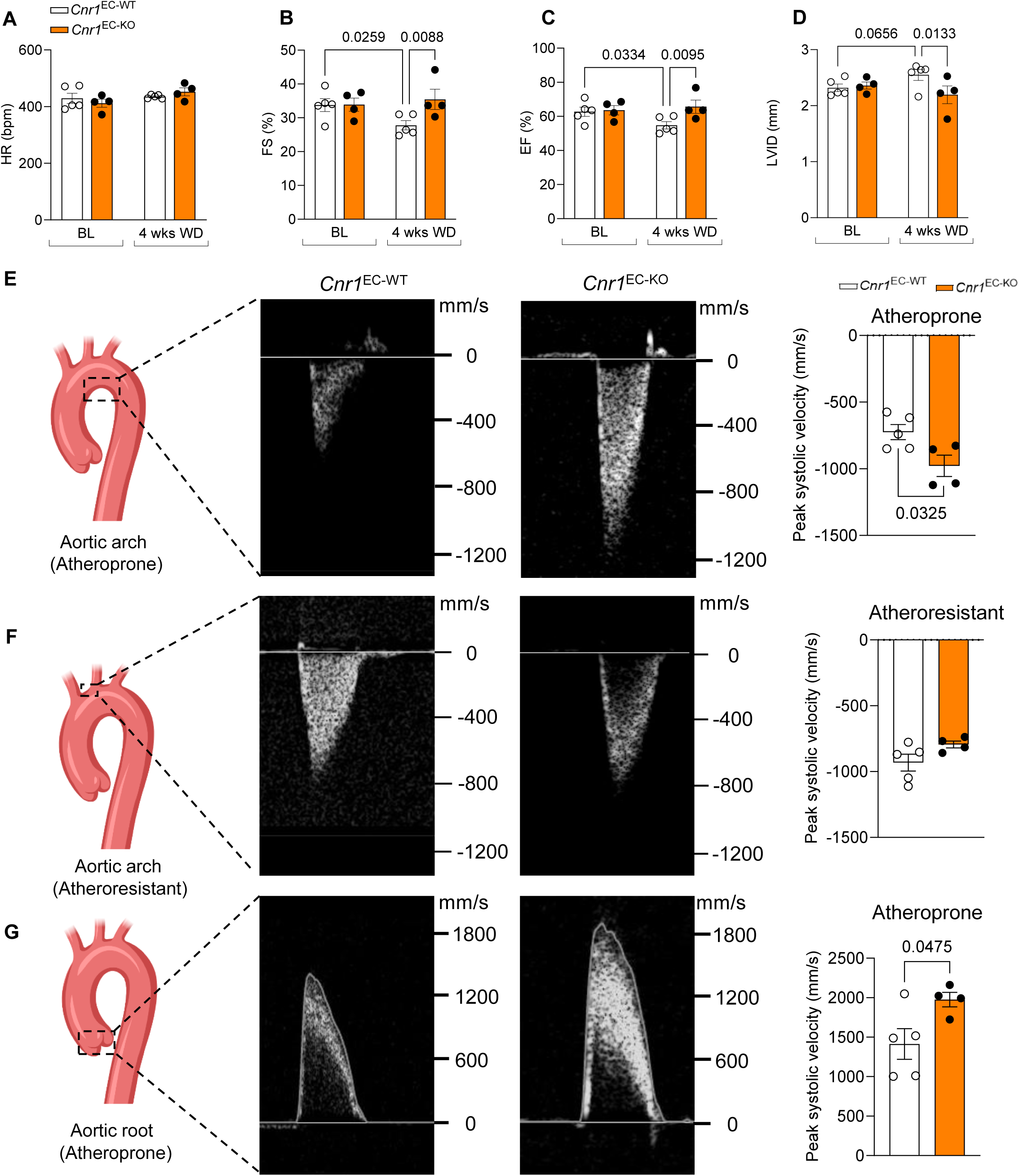
Echocardiographic assessment of cardiac function and aortic flow velocity in *Cnr1^EC-WT^* and *Cnr1^EC-KO^* mice. Age-matched female *Cnr1^EC-WT^* mice (n = 5) and *Cnr1^EC-KO^* mice (n = 4) were used to assess the cardiac function and blood flow dynamics by serial echocardiography. (**A-D**) Heart rate (HR), ejection fraction (EF), fractional shortening (FS) and end-diastolic left ventricular internal diameter (LVID) in *Cnr1^EC-WT^* and *Cnr1^EC-KO^* mice at baseline (BL) or after 4 weeks of WD. Two-way ANOVA was used to determine the significant differences. (**E-G**) Quantification of peak systolic velocity at atheroprone and atheroresistant sites of aortas at baseline. Data are shown as mean ± s.e.m. and unpaired Student’s t-test was performed.

**Supplemental Figure S4.**
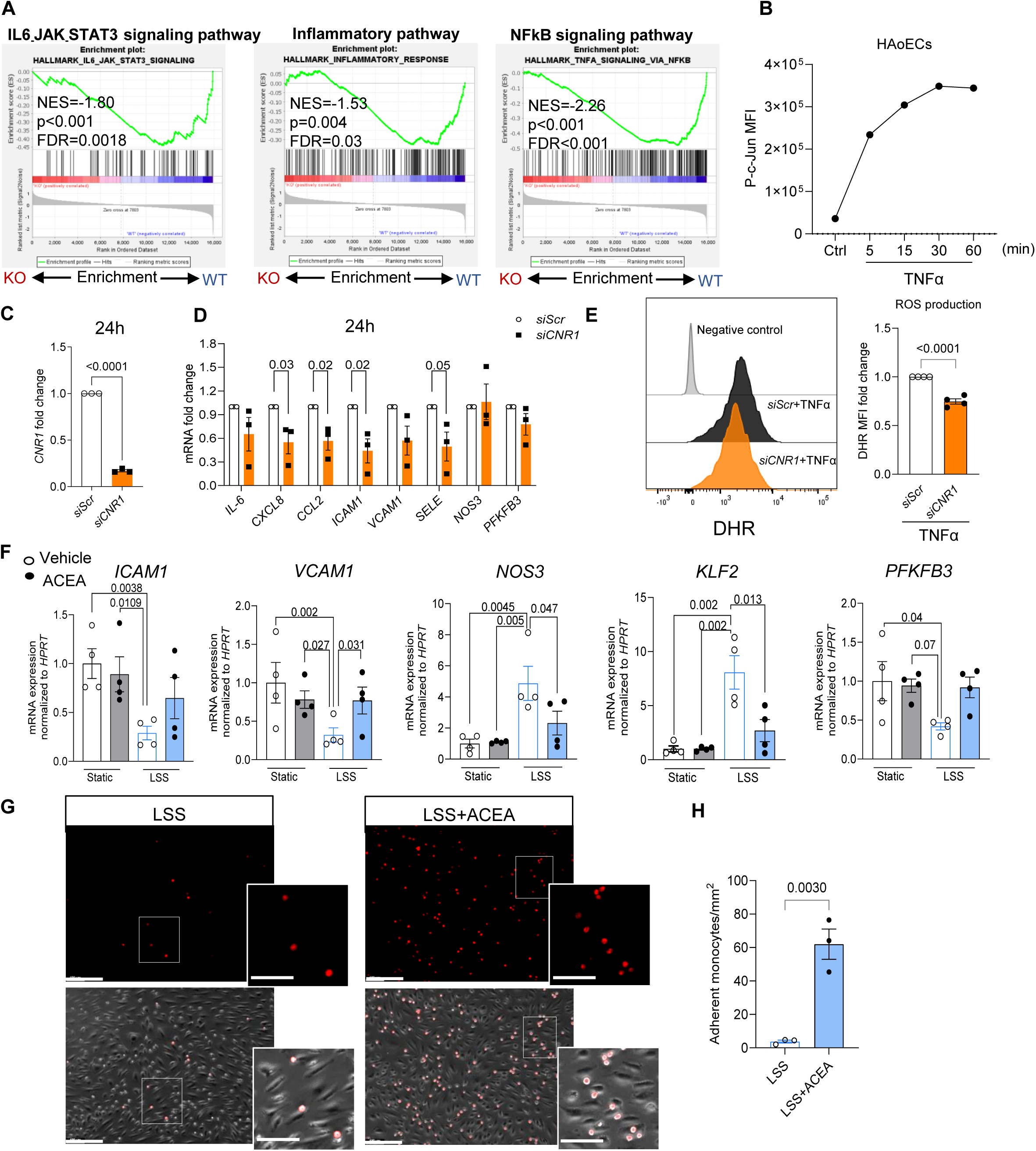
Role of endothelial CB1 in vascular inflammation and monocyte adhesion. (**A**) Pathways associated with endothelial CB1-regulated genes (GSEA). (**B**) Time course of phospho-c-Jun mean fluorescence intensity (MFI, immunostaining) in TNFα-treated HAoECs. (**C, D**) Knockdown efficiency of *CNR1* and pro-inflammatory gene expression 24h after transfection with 20 nM *CNR1* (*siCNR1*) or scrambled siRNA (*siScr*) in HAoECs (n = 3 independent experiments). (**E**) Flow cytometric analysis of ROS determined by DHR123 in HAoECs (n=4 independent experiments). (**F**) HAoECs were treated with 1 µM ACEA or vehicle under static condition or LSS (10 dye/cm^2^) for 24 h. Expression levels of shear stress related genes were assessed by RT-qPCR (n=4 independent experiments). Data were analysed by two-way ANOVA with Tukey correction for multiple comparisons. (**G**) Representative images of THP-1 monocyte adhesion (red) in vehicle- or AECA-(1 µM) treated HAoECs under LSS. Scale bar, 200 μm (overview) and 100 μm (insert). (**H**) Adherent THP-1 cells were counted in 10-15 random fields per experiment (n=3 independent experiments). Data are shown as mean ± s.e.m. and unpaired Student’s t-test (**C, D, E, H**) or two-way ANOVA with Tukey correction for multiple comparisons (**F**) was performed.

**Supplemental Figure S5.**
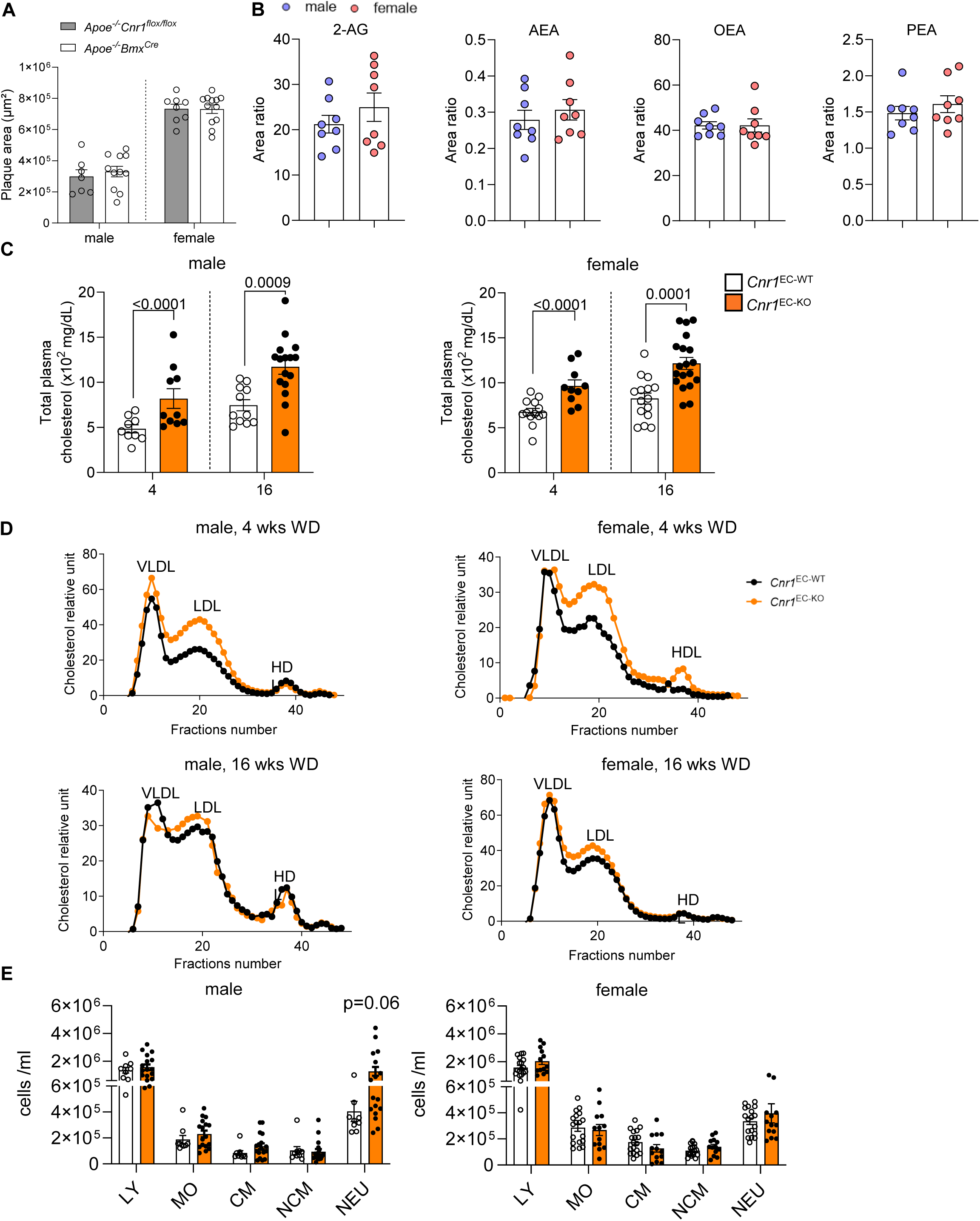
Plasma parameters and blood leukocytes in *Cnr1^EC-KO^* mice. (**A**) Quantification of the lesion area in the aortic root of *Apoe^−/−^Bmx^Cre^* and *Apoe^−/−^Cnr1^flox/flox^* mice after 16 weeks of WD (n=7-14). (**B**) Plasma endocannabinoid 2-arachidonoylglycerol (2-AG), anandamide (AEA), oleoylethanolamide (OEA), and palmitoylethanolamide (PEA) levels in male and female Apoe^−/−^ mice after 4 weeks of WD. (**C**) Plasma total cholesterol and (**D**) lipoprotein profiles in female and male *Cnr1^EC-WT^* and *Cnr1^EC-KO^* mice after 4 or 16 weeks WD (pooled plasma of 8 mice per group). (**E**) Flow cytometry analysis of lymphocytes (LY), monocytes (MO), classical (CM), non-classical monocytes (NCM), and neutrophils (NEU) in peripheral blood of *Cnr1^EC-WT^* and *Cnr1^EC-KO^* mice after 4 weeks WD. P values were obtained using an unpaired Student’s t-test.

**Supplemental Figure S6.**
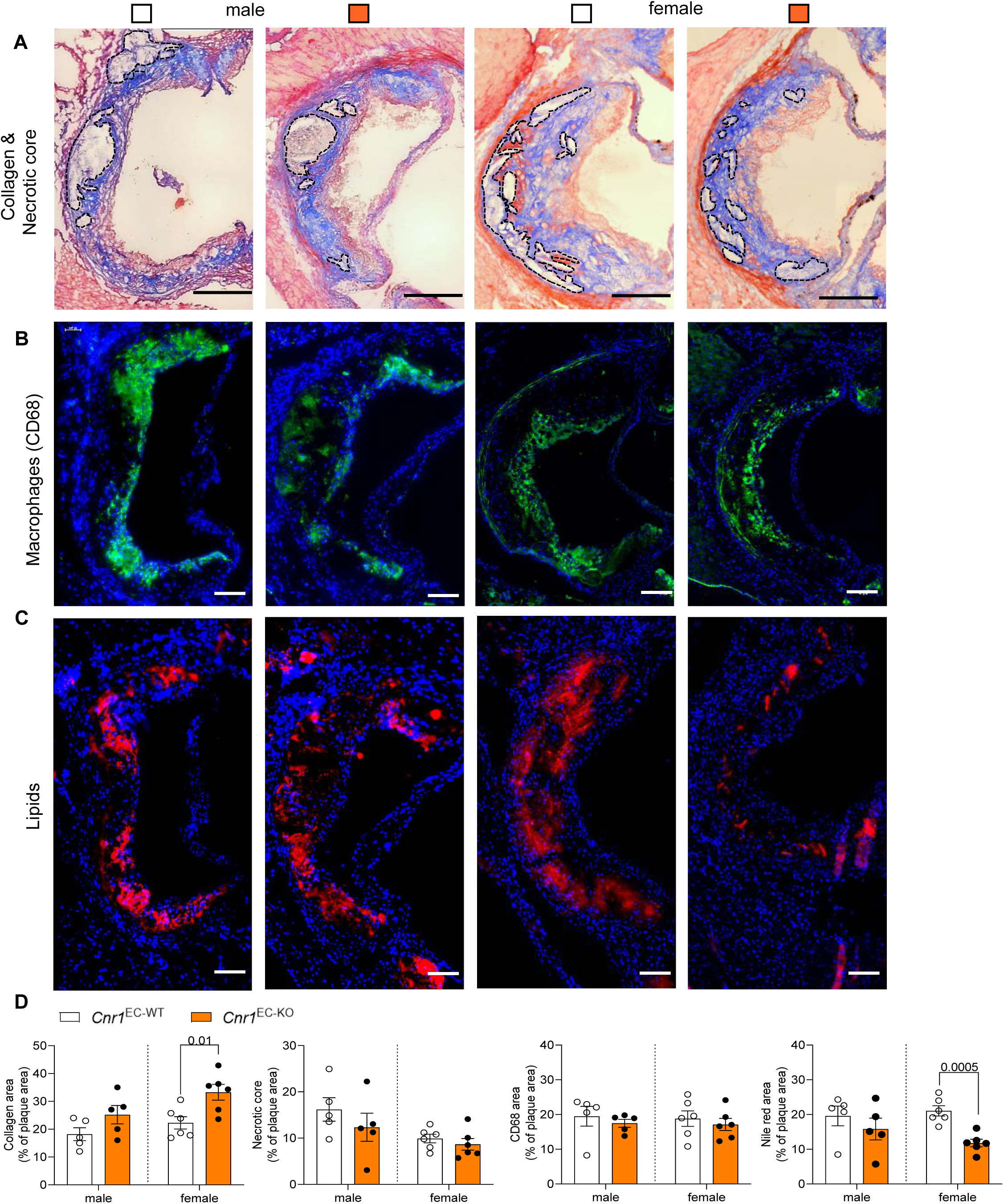
Effect of *Cnr1* deficiency on advanced plaque composition. Representative aortic root plaque images from *Cnr1^EC-WT^* and *Cnr1^EC-KO^* mice after 16 weeks WD (n=5-6) stained with (**A**) Masson’s trichrome for collagen and necrotic core area (encircled by dotted lines). Scale bar, 200 μm. (**B**) CD68 immunostaining for macrophages and (**C**) Nile red staining for intracellular lipids. Scale bar, 100 μm. (**D**) Quantification of relative areas per plaque area. Data are shown as mean ± s.e.m.; unpaired Student’s t-test was conducted separately for males and females.

**Supplemental Figure S7.**
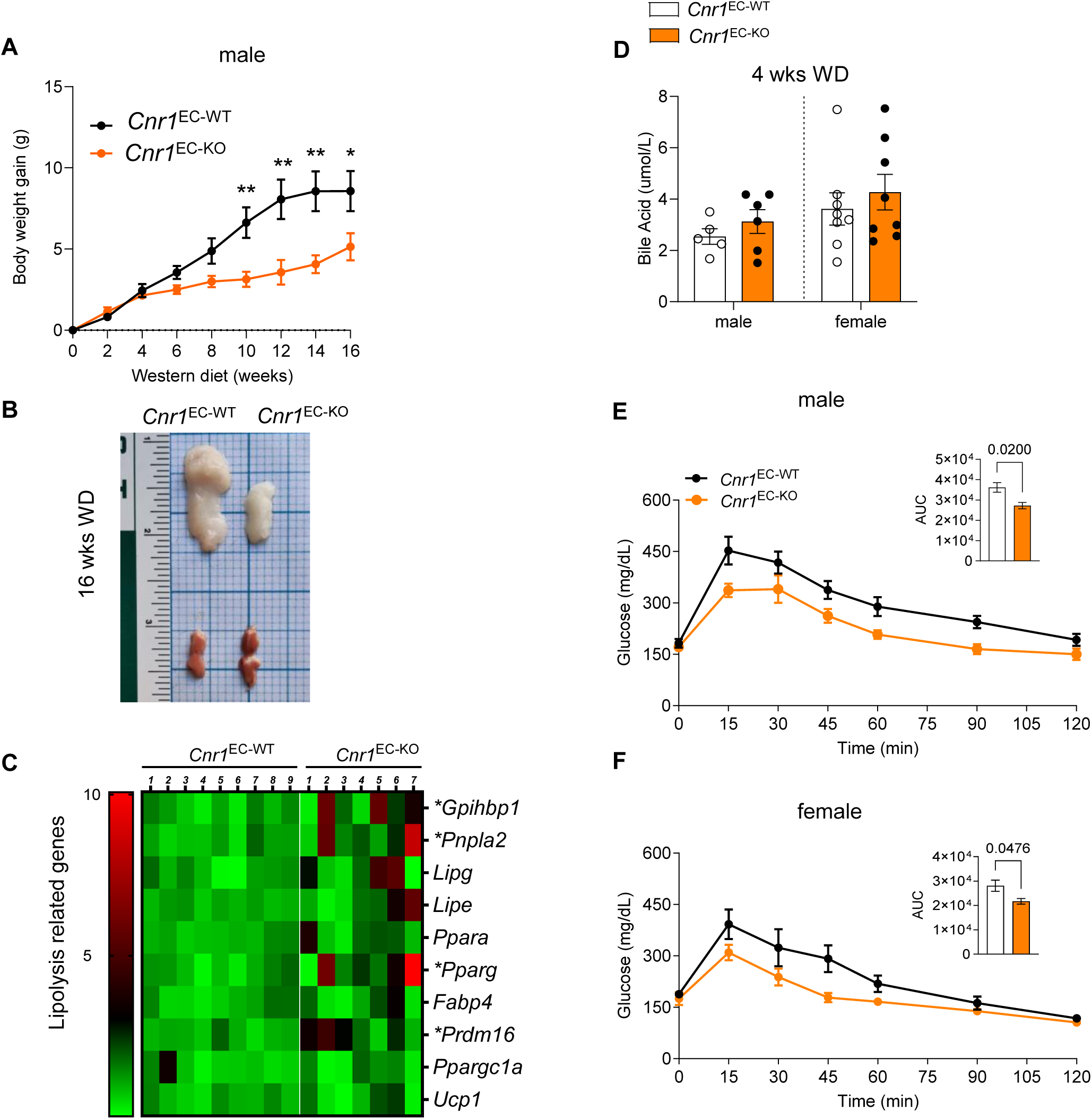
Impact of endothelial *Cnr1* deficiency on metabolic parameters in males. (**A**) Body weight gain over 16 weeks WD in male *Cnr1^EC-WT^* and *Cnr1^EC-KO^* mice (n=7-8). (**B**) Representative images of epididymal white (eWAT) and brown adipose tissue (BAT) from male mice after 16 weeks of WD. (**C**) Gene expression analysis (RT-qPCR) in BAT of male *Cnr1^EC-WT^* and *Cnr1^EC-KO^* mice after 4 weeks of WD (n = 9). (**D**) Plasma bile acid levels in male (n=5-6) and female (n=8) *Cnr1^EC-WT^* and *Cnr1^EC-KO^* mice after 4 weeks of WD. (**E-F**) Plasma glucose levels during intraperitoneal glucose tolerance test and area under the curve (AUC) in *Cnr1^EC-WT^* and *Cnr1^EC-KO^* after 4 weeks of WD (n = 4). Data are shown as mean ± s.e.m. and two-tailed Student’s t-test was performed. **p<0.01; *p<0.05

**Supplemental Figure S8.**
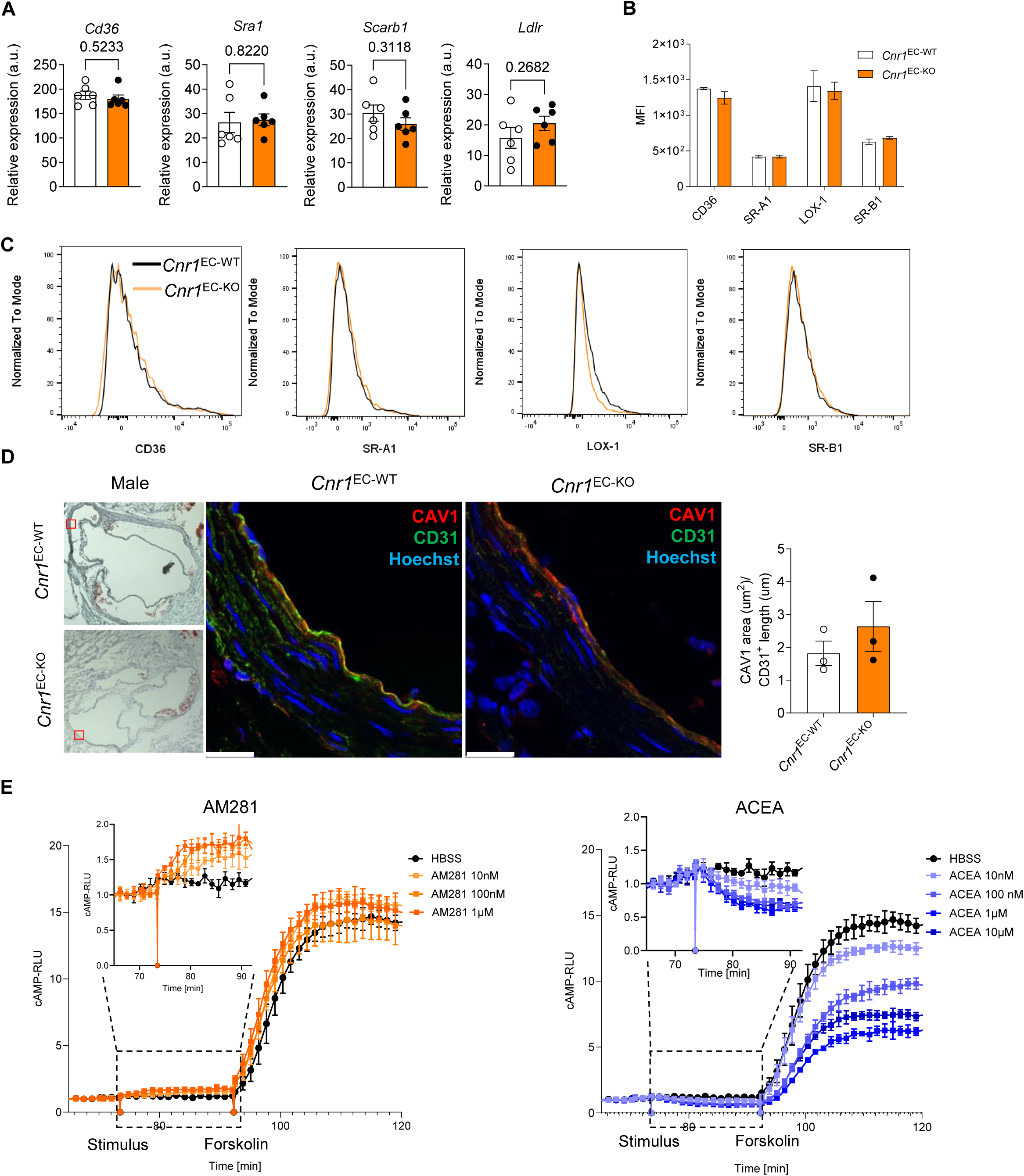
Impact of endothelial *Cnr1* deficiency on endothelial lipid receptors and CB1-dependent regulation of cAMP. (**A**) Expression of lipid receptors retrieved from RNA sequencing data of sorted aortic ECs from female *Cnr1^EC-WT^* and *Cnr1^EC-KO^* mice (n=6). (**B**, **C**) Flow cytometric analysis of lipid receptor surface expression on aortic endothelial cells of female *Cnr1^EC-WT^* and *Cnr1^EC-KO^* mice after 4 weeks WD; gated as live CD45^−^CD31^+^CD107a^+^ (n=2-4). (**D**) Representative confocal images of immunostainings and quantification (n=3) of caveolin-1 (CAV1, Red), CD31 (green) and Hoechst33342 (nuclei, blue) in aortic root sections of male *Cnr1^EC-WT^* and *Cnr1^EC-KO^* mice after 4 weeks WD. Scale bar, 20 μm. (**E**) Glosensor cAMP reporter assay in *CNR1* and cAMP-luciferase expressing HEK293 cells, treated with buffer (HBSS) alone or CB1 antagonist AM281 or agonist ACEA, followed by addition of the adenylate cyclase activator forskolin. Data are shown as mean ± s.e.m. and unpaired Student’s t-test was performed.

**Supplemental Figure S9.**
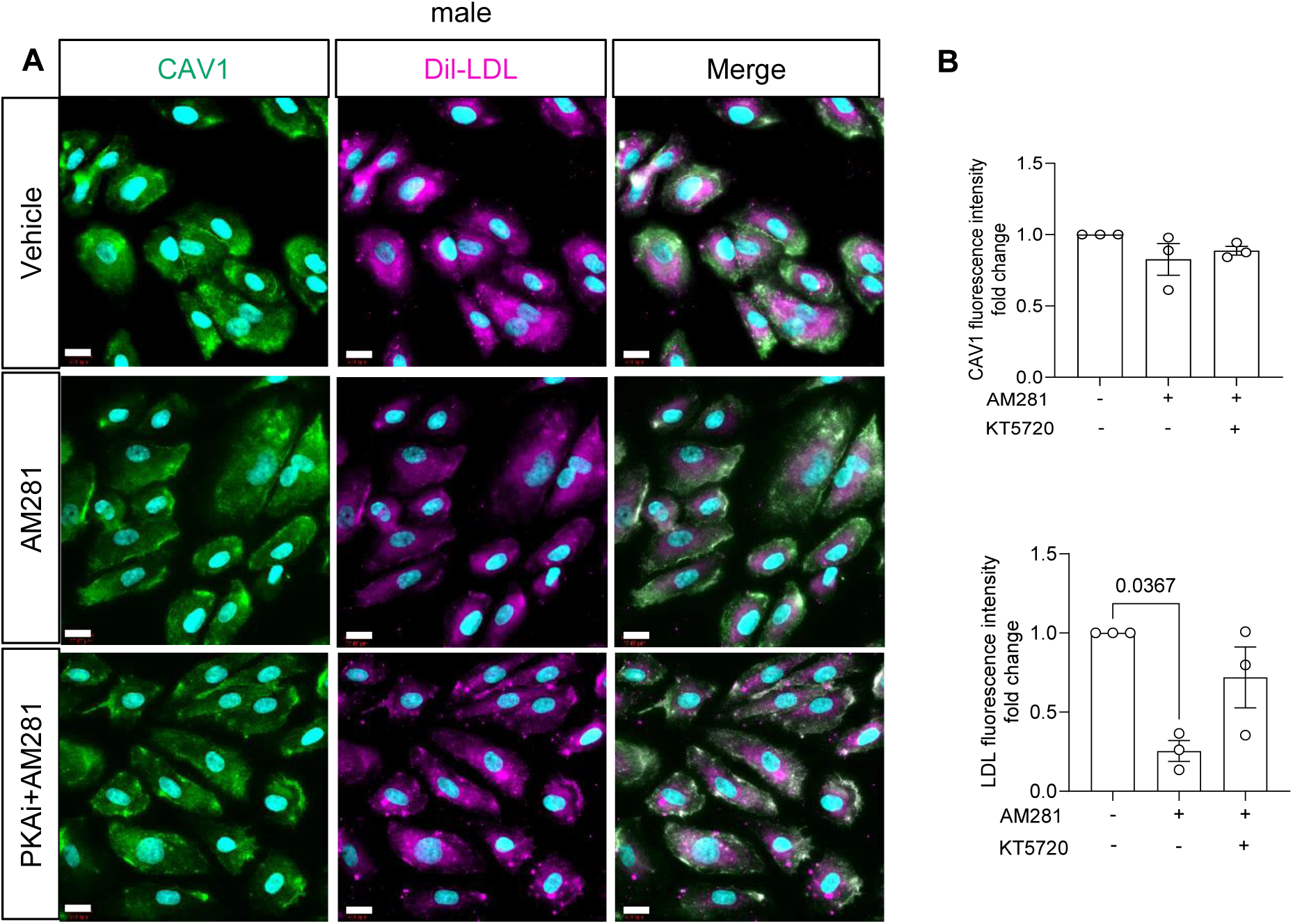
DiI-LDL uptake in HAoECs of male donors. (**A**) Representative immunofluorescence analysis of CAV1 expression and Dil-LDL uptake in HAoECs of male donors treated with 1 μM AM281 alone or in the presence of 1 μM PKA inhibitor (KT5720) or vehicle (DMSO) under OSS for 24 h. Scale bar, 20 μm, and (**B**) quantification of n=5 independent experiments. Data are shown as mean ± s.e.m. and one-way ANOVA with Tukey correction was performed.

**Supplemental Figure S10.**
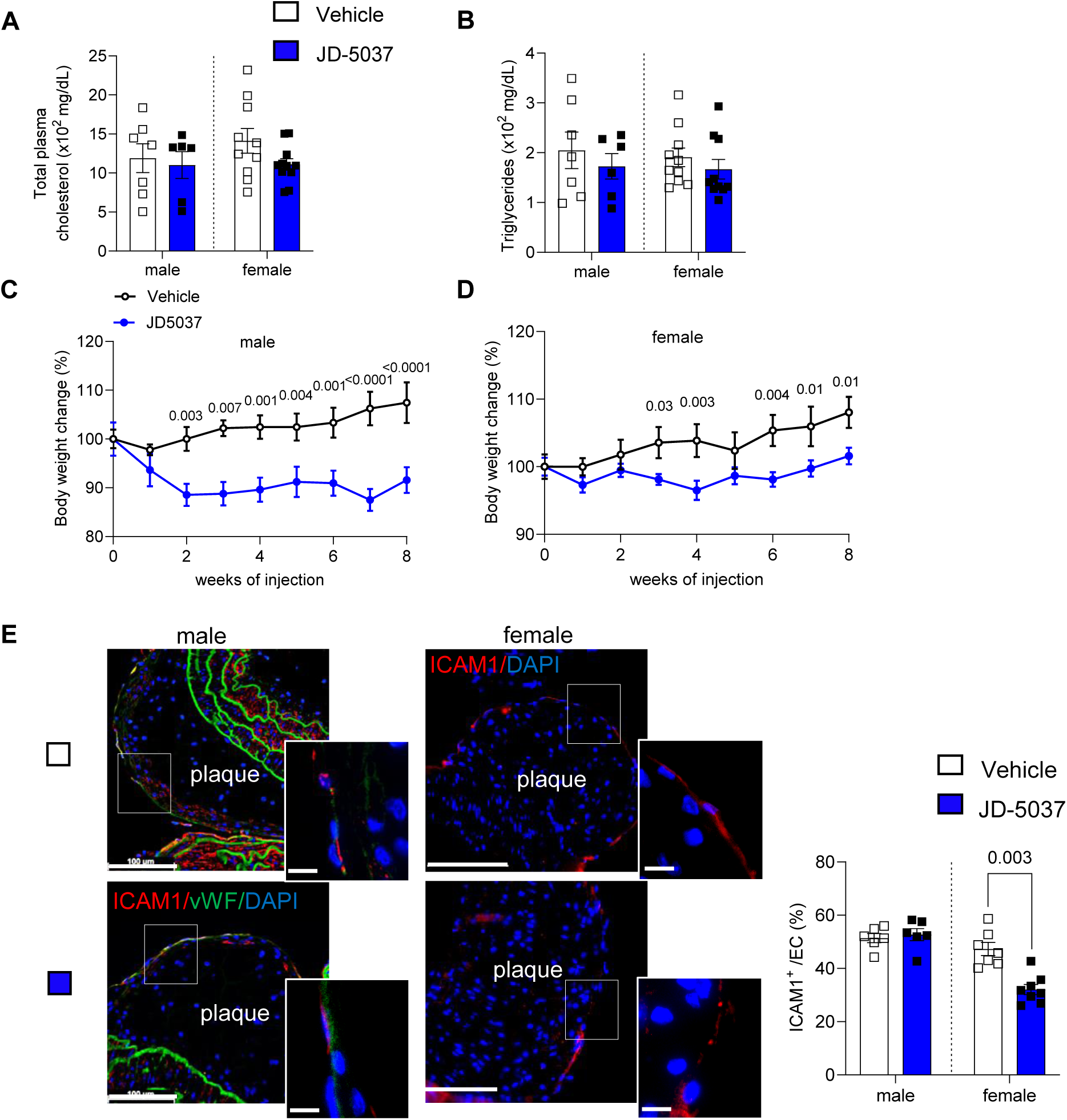
Impact of peripheral CB1 antagonist JD5037 on metabolic parameters and plaque endothelial ICAM1 expression. (**A**) Plasma cholesterol and (**B**) triglyceride levels in male and female *Ldlr^−/−^* mice (n=6-8) after 16 weeks WD and daily JD or vehicle injections for the last 8 weeks. Body weight changes in male (**C**) and female (**D**) *Ldlr^−/−^* mice (n=6-8) during the 8 weeks of JD5037 or vehicle treatment. (**E**) Representative images and quantification of ICAM1 (red) positive endothelial cells (van Willebrand factor, vWF; green) in aortic arch lesions of male and female *Ldlr^−/−^* mice (n=6-8) treated with vehicle or JD5037. Nuclei were stained with DAPI (blue). Scale bar, 100 μm (overview) and 10 μm (insert). Each data point represents a mouse, and all data are expressed as mean ± s.e.m. Unpaired Student’s t-test was performed, and male and female data were analysed independently.

**Supplemental Figure S11.**
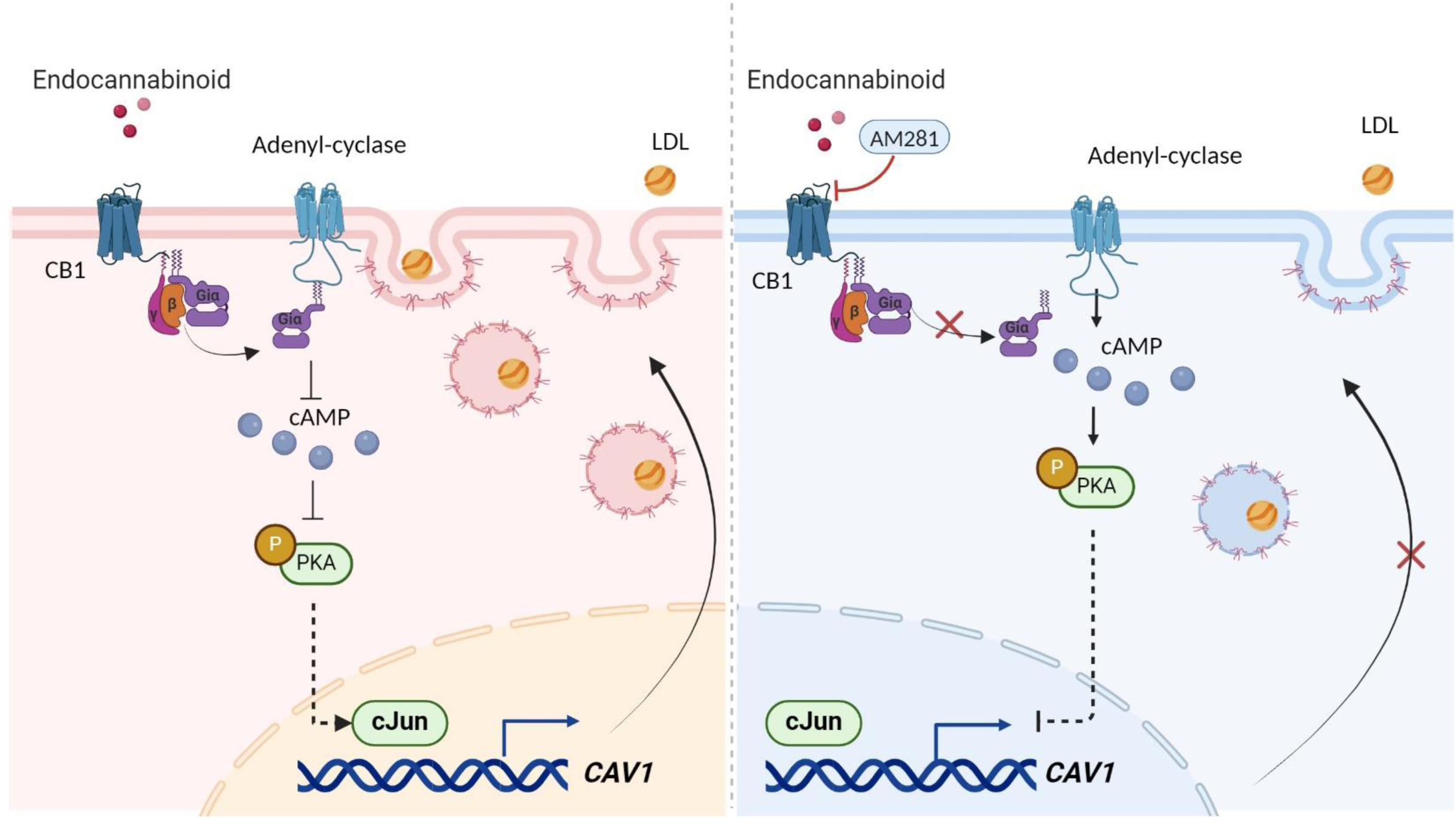
Proposed mechanism how CB1 regulates endothelial CAV1 expression and LDL uptake. (**Left**) Endocannabinoid-CB1 signaling leads to a G_i_-dependent inhibition of adenylate cyclase and consequently to an inhibition of intracellular cAMP production. Reduced cAMP-dependent PKA activity prevents PKA-dependent inhibition of *CAV1* gene expression and LDL transcytosis. (**Right**) Antagonism of CB1 prevents the depletion of intracellular cAMP and thereby enables PKA activation, which inhibits *CAV1* gene expression and caveolae-dependent LDL uptake.

## SUPPLEMENTAL METHODS

### Animal model of atherosclerosis

To generate mice with endothelial *Cnr1* deficiency on atherogenic background, *Cnr1^flox/flox^* mice (kindly provided by Beat Lutz)^1^ were first crossed with *Apoe^−/−^* mice to generate *Apoe^−/−^ Cnr1^flox/flox^* mice*. Apoe^−/−^Cnr1^flox/flox^* mice were then crossed with *Bmx^CreERT^* mice^2^ to obtain *Apoe^−/−^Bmx^Cre(+/-)^Cnr1^flox/flox^*mice (referred to as *Cnr1^EC-KO^*). The deletion was induced by intraperitoneal injection (i.p.) of tamoxifen (1 mg per 20g body weight, dissolved in corn oil) at 8 weeks of age, administered for 5 consecutive days to induce *Bmx^CreERT^* transgene expression for selective Cre recombination in arterial ECs. Tamoxifen was also injected into *Apoe^−/−^ Bmx^Cre(+/-)^*and *Apoe^−/−^Cnr1^flox/flox^* control mice. Following tamoxifen induction, the mice rested for 10 days before baseline harvest or starting a Western diet (WD) consisting of 21% fat and 0.2% cholesterol (Ssniff, TD88137) for either 4 or 16 weeks. In a separate set of experiments, *Ldlr^−/−^* mice^3^ were first fed 8 weeks of WD to induce atherosclerotic plaque formation. Subsequently, the mice were randomly divided into two groups to receive daily i.p. injections of either JD5037 (3 mg/kg) or vehicle (10% DMSO, 40% PEG300, 5% Tween 80, 45% saline) with continuous WD feeding for a total duration of 16 weeks. At the study endpoints, mice were anesthetized with ketamine/xylazine, and blood was obtained via cardiac puncture. Heart, aorta, adipose tissue, and liver were harvested after PBS perfusion. Animals were housed in ventilated cages, with 4 to 6 mice per cage. The environment was air-conditioned, with a 12-hour light-dark cycle and a temperature of 23°C and 60% relative humidity. All animal procedures were approved by the local Ethics committee (District Government of Upper Bavaria; license number: 55.2-1-54-2532-111-13 and 55.2-2532.Vet_02-18-114) and conducted in accordance with the institutional and national guidelines and following the ARRIVE guidelines.

### Permeability assay

Evans blue solution (0.5 %) was prepared in saline and sterilized by filtering. Mice were injected into the tail vein with 200 µl of Evans blue solution and euthanized 30 min post-injection as described above. The entire aorta was collected, fixed with 4 % paraformaldehyde (PFA) for 30 min, and positioned on slides for imaging (image settings for Evans blue excitation peaks: 470 nm and 540 nm, with an emission peak at 680 nm). Tilescan z-stacks of the whole aorta were taken with Leica DM6000B microscopes and images analyzed with Leica Application Suite LAS V4.3 software.

### Plasma and liver total cholesterol measurement

Total plasma cholesterol concentrations were measured with a colorimetric assay (CHOD-PAP; Roche) and microplate reader (Infinite F200 PRO, Tecan). Plasma from WD-fed mice was diluted at a 1:9 ratio with 0.9% saline for analysis. Liver tissue (50-70 mg) was homogenized in 500 μl of 0.1% NP-40 in PBS using a Tissue Lyser bead mill (Qiagen) and then centrifuged to remove insoluble material. The supernatant was collected and diluted in 0.1% NP-40 PBS. Liver total cholesterol was normalized to the protein concentration, which was determined via a bicinchoninic acid (BCA) assay (Bio-Rad, USA).

### Lipoprotein profile analysis

For lipoprotein separation, plasma samples from 8 mice per group were pooled (0.2 ml) and subjected to fast performance liquid chromatography (FPLC) gel filtration on two Superose 6 columns connected in series as described previously.^4^

### Endocannabinoid measurement

Lipid extraction from mouse plasma was performed on ice. Samples were thawed on ice and spiked with 10 μL internal standard mix. Subsequently, 100 μL ammonium acetate buffer (0.2 M, pH 4) were added. After extraction with 1 mL methyl tert-butyl ether (MTBE), tubes were thoroughly mixed for 4 min using a bullet blender blue (Next advance Inc., Averill Park, NY, USA) at speed 6, followed by a centrifugation step (16,000×g, 10 min, 4 °C). Next, 950 μL of the upper MTBE layer was transferred into a clean 1.5 mL Safe-Lock Eppendorf tube. Samples were dried in a speedvac (Eppendorf, 45 min, 30 °C) and reconstituted in acetonitrile/methanol (50 μL, 70:30, v/v). The samples were thoroughly mixed for 15 min, followed by a centrifugation step (16,000×g, 4 min, 4 °C) and transferred to an LC–MS vial (9 mm, 1.5 mL, amber screw vial, KG 090188, Screening Devices) with insert (0.1 mL, tear drop with plastic spring, ME 060232, Screening devices). 5 μL was injected into the LC–MS/MS system. A targeted method covering endocannabinoids and related N-acylethanolamines (NAEs) with slight modifications was applied.^5^ A QTRAP 6500+ (AB Sciex, Concord, ON, Canada) coupled to an Exion LC AD (AB Sciex, Concord, ON, Canada). MS/MS experiments were done with a Turbo V source (AB Sciex, Concord, ON, Canada) operated with ESI probe. The separation was performed in a BEH C8 column (50 mm × 2.1 mm, 1.7 μm) from Waters Technologies (Mildford, MA, USA) maintained at 40°C, with the flow rate at 0.4 mL/min. The mobile phase was consisted of 2 mM HCOONH4, 10 mM formic acid in water (A), ACN (B), IPA (C). The gradient was the following: starting conditions 20% B and 20% C; increase of B from 20% to 40% between 1 min and 2 min; maintaining B at 40% and C at 20% between 2 min and 7 min; increase of C from 20% to 50% between 7 min and 8 min; maintaining B at 40% and C at 50% between 8 min and 10 min; returning to initial conditions at 10.5 min and re-equilibration for 1.5 min. The triple quadrupole mass spectrometer operated in polarity switching mode and all analytes were monitored in dMRM mode. Data was acquired using Sciex OS Software V2.0.0.45330 (AB Sciex). Assigned MRM peaks from the acquired data were integrated using SCIEX OS (version 2.1.6) Software and signals were corrected using proper internal standards. Blank effects for each analyte were checked by comparing proc blank samples to quality control (QC) samples. The precision and reproducibility of the analytical process were checked using the relative standard deviations (RSDs) of the QCs.

### Glucose tolerance test

Mice underwent a 6-hour fasting period with unrestricted access to water prior to i.p. injection of glucose (2 g/kg). Blood samples were collected from the caudal vein to measure plasma glucose levels at specific time intervals (0, 15, 30, 60, and 120 min) using a glucometer (Accu-Chek, Mannheim, Germany).

### Serial echocardiographic assessment

Transthoracic echocardiography was performed with the Vevo® 3100 Imaging System (FUJIFILM VisualSonics; Toronto, Canada) using the MX550 transducer (25-55 MHz). Mice were initially anesthetized with 4% isoflurane supplemented with oxygen, which was reduced to 2-3% during image acquisition. Peak aortic velocities were obtained from the color Doppler-mode aortic arch view. Systolic and diastolic cardiac functions were analyzed in M-mode of the left ventricular parasternal long-axis view. Image analysis and calculations were done using the VevoLAB Version 5.7.0 (FUJIFILM VisualSonics, Toronto, Canada).

### Histology and immunofluorescence of atherosclerotic plaques

Atherosclerotic lesion sizes were analyzed in aortic root cryosections, aortic arch paraffin sections and *en face* prepared aortas. Mouse hearts were isolated after perfusion with PBS and embedded in Tissue-Tek O.C.T. compound (Sakura) and frozen for cutting into 5 µm cross-sections. Lesion size within aortic roots was quantified after Oil-Red-O (ORO) staining using 8 sections per heart, separated by 50 μm from each other. Aortic arches were fixed overnight in 1% paraformaldehyde (PFA) and embedded in paraffin for longitudinal sectioning (4 µm). Lesion size was quantified after H&E staining, and the average plaque size was calculated from 3-4 sections per arch separated by 40 μm from each other. For *en face* analysis of plaques in the aortic arch and descending aortas, the vessels were fixed overnight in 1% PFA and carefully opened and pinned on black rubber plates for imaging.

Aortic root cryosections were also used for plaque composition analysis, using 3-4 sections per mouse heart for quantification. For assessing macrophage content, acetone-fixed sections were incubated overnight with an antibody against CD68 at 4 °C, followed by anti-rat-AF488 for 1 hour at room temperature (Supplemental Tables 1-2). Lipid droplets were stained with Nile Red (N3013, Sigma-Aldrich) for 5 minutes, followed by nuclear -staining with Hoechst 33342 for 5 minutes. Plaque collagen and necrotic core content were assessed by Masson’s trichrome staining (stain kit Sigma HT15) in accordance with the guidelines for experimental atherosclerosis studies by the AHA.^6^ For CAV1 detection, aortic root cryosections were fixed with 4% PFA, permeabilized with 0.1% triton X-100 in PBS, blocked for 1h and then incubated overnight with anti-CAV1 and anti-CD31 antibodies at 4°C, followed by anti-rat-AF488 and anti-rabbit-AF647 for 1 hour at room temperature. Corresponding isotypes were included as negative staining controls (Supplemental Table 3). Nuclei were stained with Hoechst 33342 for 5 min. Images for CAV1 detection were taken with a Leica SP8 3 X confocal microscope (Leica) and images were digitized with constant exposure time, gain and offset. Results were expressed as positive staining area (µm^2^) normalized to the length of the endothelial cell layer (µm) analyzed with the Leica Application Suite LAS V4.3 software. Images of *en face* prepared vessels for plaque quantification were taken with a Leica M205 FCA microscope equipped with a 2.5x objective. All other images were taken with a fluorescence microscope (DM6000B) connected to a monochrome digital camera (DFC365FX, Leica) or connected to a bright field digital camera (DMC6200, Leica) equipped with Thunder technology for computational clearing and analyzed with the Leica Application Suite LAS V4.3 software. All plaque data are expressed as average per section and mouse heart.

### ICAM1, VCAM1 and CAV1 staining in aortic arch

Aortic arches were fixed in 1% PFA overnight and embedded in paraffin for longitudinal sectioning (5 µm). The tissue sections were first deparaffinized and subjected to antigen retrieval. Tissue sections were then incubated overnight at 4°C with primary antibodies targeting CAV1, ICAM1, and vWF, or VCAM1 and vWF (Supplemental Tables 1-3) after 30 min blocking at RT, followed by corresponding Alexa Fluor secondary antibodies for 30 min at RT. Slides were mounted with Vectashield mounting media with DAPI. Quantification was conducted on 4 sections per animal, with a 50 µm interval between each section imaged by a Leica Thunder DM6000B microscope at 20x magnification. To determine the percentage of ICAM1/VCAM1/CAV1-positive ECs, the number of ICAM1/VCAM1/CAV1-positive cells was calculated and normalized to endothelial cell numbers using LAS V4.3 software (Leica).

### Immunofluorescence staining of brown adipose tissue

Brown adipose tissue (BAT) was fixed overnight in 4% PFA and embedded in paraffin for longitudinal sectioning (4 µm). The paraffin sections were first deparaffinized and subjected to antigen retrieval. Tissue sections were then incubated overnight at 4°C with primary antibody against GPIHBP1 and vWF after 30 min blocking at RT, followed by Cy3 donkey anti-rabbit and Cy5 anti-sheep secondary antibody staining for 30 min at RT (Supplemental Tables 1-3). Slides were mounted with Vectashield mounting media with DAPI. Images were taken with a Leica Thunder DM6000B microscope (Leica) and quantified using LAS V4.3 software (Leica). Five images were captured per section, and 3-4 sections per mouse were quantified, with the results presented as the mean value per animal.

### *En face* immunofluorescence staining of thoracic aortas

Prior to harvest, the thoracic aortas were perfused with 20ml pre-cooled PBS containing 20% FBS and 4% PFA. Subsequently, the vessels were carefully opened and transferred to a 12-well plate, containing 4% PFA for 20 min fixation. Whole-mount immunofluorescence staining was performed based on a published protocol.^7^ The vessels were permeabilized with 0.1% Triton X-100 in PBS, washed, and blocked with PBS containing 1% horse serum and 1% BSA for 1 h at room temperature. Aortas were incubated overnight with primary antibodies against CD144 and ICAM1 on a rocking platform at 4°C, followed by 1 h incubation with donkey anti-rat and goat anti Armenian hamster Alexa Fluor secondary antibodies (Supplemental Tables 1-3). Aortas were mounted with Vectashield mounting media with DAPI and imaged using a confocal microscope (TCS-SP5, Leica) at 488 and 550 nm, respectively. Endothelial cells were identified as CD144 positive under the same setting. Atheroprone and atheroprotective regions were chosen for high magnification imaging after low magnification tile scanning of the entire aorta. The mean fluorescence intensity (MFI) of maximum projections of images was analyzed using LAS X Office image processing software and ImageJ.

### Fluorescence *in situ* hybridization

Endothelial Cnr1 detection was performed by *in situ* hybridization combined with immunostaining employing viewRNA cell plus kit (Thermo Fisher).^8^ Arch and thoracic aortae *en face* prepared thoracic aortas of *Apoe^−/−^* mice were fixed in Paxgene for 1h and 30 min and en face prepared for tissue stabilizing in a stabilized solution overnight at 4°C. The entire procedure was performed in RNase-free conditions and using an RNase inhibitor cocktail alongside the hybridization steps. Tissues were incubated with a custom probe designed for murine *Cnr1* (VB6-17606, Affymetrix) and incubated at 40±1°C for 2 h for the target probe hybridization process. The hybridization probe was diluted in probe set diluent to achieve a final concentration of 5 μg/ml. Subsequently, the tissues were washed and incubated with the pre-amplifier mixture at 40±1°C for 90 min, and then exposed to the amplifier mixture for an additional 1 h to enhance the signal. After the hybridization process, tissues were incubated with approprioate fluorescently labelled probes (Type 6) for 1 h. Tissues underwent a thorough wash followed by 30 min of incubation with a fixation/permeabilization buffer, another wash, and a 1-h incubation with blocking buffer at room temperature. Subsequently, they were stained with anti-CD31 antibody overnight at 4°C in a humid chamber, followed by a 1h incubation with AlexaFluor488 anti-rat secondary antibody at room temperature. Tissues were transferred to glass slides after final washing and mounted with Vectashield mounting media with DAPI.

Aortic root cryosections were collected on RNAse-free slides (pre-treated with RNAse ZAP). The cryosections were fixed in 4% PFA for 5 min, followed by treatment with pre-warmed 10 μg/ml proteinase K (diluted in PBS) for 5 min at RT. Subsequently, post-fixation was carried out with 100% ethanol for 1 min. The prepared slides were then mounted with SecureSeal™ hybridization chambers. The slides were then placed on the in-situ adapter of an Eppendorf Mastercycler machine and incubated at 40±1°C following the same hybridization procedure as described above. Afterwards, the chambers were removed, washed and stained with anti-CD31 and Hoechst 33342 as described above. Images were taken using a confocal microscope TCS-SP5. An average of 5-10 images per section and per vessel were acquired and subsequently quantified using ImageJ software.

### *Ex vivo* imaging of lipid uptake in perfused carotid arteries

Endothelial Dil-LDL (3,3’-dioctadecylindocine-low density lipoprotein) uptake was assessed in murine carotid arteries of *Cnr1^EC-WT^*and *Cnr1^EC-KO^* mice (n=5-7) mounted in perfusion chambers as previously described.^9,10^ The vessels were first incubated with anti-CD31 (Supplemental Table 4) under a static pressure of 80 mmHg for 10 min at 37°C. After washing, DiL-LDL was loaded onto the arteries and incubated for 90 min under 80 mmHg at 37°C. Following the removal of unbound antibodies and DiL-LDL through artery flushing, tissues were imaged using a Leica SP5 IIMP two-photon laser scanning microscope coupled to a Ti:sapphire laser (Spectra Physics MaiTai DeepSee) tuned at 800 nm. A 20× NA1.00 (Leica) water dipping objective was utilized, and spectral detection employed internal Hybrid Diode detectors tuned for optimal contrast between various targets while maintaining sufficient fluorescence signal intensity from the arterial wall. Three-dimensional image processing and quantification of DiL-LDL distribution per endothelial cell were performed using Leica LASX 3.11 software, utilizing 3D analyser and lightning plugins.

### Flow cytometry analysis of blood leukocytes

Freshly collected whole blood (50 µl) was transferred to ice-cold FACS tubes and subjected to red blood cell lysis with ammonium-chloride-potassium (NH_4_Cl (8,024 mg/l), KHCO_3_ (1,001 mg/l), EDTA.Na_2_·2H_2_O (3,722 mg/l)) buffer for 10 min at RT. Cell suspensions were incubated with an antibody mix (Supplemental Table 5) for 30 min at 4°C in the dark. Then, the cell suspensions were washed with 1 ml of FACS buffer and then centrifuged into 300-500 µl FACS buffer based on the cell count. The cells were washed and acquired with a BD FACSCanto II flow cytometer (BD Biosciences) and analysed with FlowJo v10.2 software (Tree Star, Inc). Cells were gated as singlets, live and CD45^+^CD11b^+^ myeloid subsets and further gated as CD115^+^Ly6G^−^ (monocytes) and CD115^−^Ly6G^+^ (neutrophils).

### Flow cytometry sorting of aortic and adipose tissue endothelial cells

Murine aortas spanning from the aortic arch to the iliac bifurcation were isolated after perfusion with PBS and digested with collagenase IV and DNase I (Supplemental Table 6) at 37°C for 40 min. The interscapular brown adipose tissue (BAT) was collected, cut into small pieces and digested with collagenase I, collagenase XI, DNase I, and hyaluronuclease (Supplemental Table 7) at 37°C for 30 min. Subsequently, the digested tissues were washed and filtered through a 30-μm cell strainer (Cell-Trics, Partec). The resulting single-cell suspensions were stained with antibody cocktails (Supplemental Table 5) and sorted using a BD FACS Aria III Cell Sorter (BD Biosciences), gated as live CD45^low^CD31^high^CD107a^high^ aortic endothelial cells or live CD45^−^CD31^+^ as BAT endothelial cells, respectively. The sorted cells were deep-frozen in 2× TCL buffer (Qiagen) plus 1% b-mercaptoethanol at −80°C until subsequent RNA extraction and library preparation.

### RNA sequencing

RNA sequencing of 10.000 sorted aortic and BAT ECs isolated from *Cnr1^EC-WT^* or *Cnr1^EC-KO^* mice after 4 weeks WD (n=6) was performed using the prime-seq protocol that can be found on protocols.io (dx.doi.org/10.17504/protocols.io.s9veh66).^11^ For differential gene expression analysis, DESeq2 (Version 1.37.4), a Bioconductor package implemented in R version 4.2.0 (2022-04-22), was utilized. The analysis was executed on an Ubuntu 20.04.3 LTS system, employing the negative binomial distribution for the necessary computations. Initially, size factors and sample dispersion were estimated, followed by utilizing Negative Binomial GLM fitting and computing Wald statistics using DESeq2. DESeq2 was applied to identify differentially expressed genes (DEGs) based on the criterion of an adjusted P value below 0.10.^12,13^ The volcano plot was created using the ggplot2 package (Version 3.4.0). Gene ontology (GO) enrichment analysis was conducted using the enrichplot (Version 1.17.2) Bioconductor package,^14^ employing DEGs filtered by an adjusted P value < 0.10. The CHEA3 web server was employed to predict the top 15 transcription factors (TFs) regulating DEGs.^15^ Gene Set Enrichment Analysis (GSEA) was performed using GSEA software (version 4.3.2), which was tailored for the Windows operating system. The analysis utilized curated gene sets (M2), ontology gene sets (M5) and mouse-ortholog hallmark gene sets sourced from mouse collections.^16^ Pathways were considered significant according to predefined criteria, encompassing a normalized enrichment score below −1 or above 1, a false discovery rate below 0.25, and a nominal P value less than 0.05.^17^ The bulk RNA sequencing dataset is accessible under GEO accession number GSE260826.

### Quantitative real time PCR

Total isolated RNA from cells (RNeasy Plus Mini Kit, Qiagen) or tissues (peqGOLD, 13-6834-02, VWR Life Science) was reverse transcribed (PrimeScript RT reagent kit, TaKaRa) to cDNA. Real-time qPCR was performed with the QuantStudio™ 6 Pro Real-Time PCR System (ThermoFisher) using the GoTaq Probe qPCR Master Mix (Promega). Primers and probes were purchased from Life Technologies (Supplemental Tables 8-9). Hypoxanthine-guanine-phosphoribosyltransferase (*Hprt*) was used as the endogenous control. For BAT samples, ubiquitin C (*Ubc*), was utilized as endogenous reference. Target gene expression was normalized to the endogenous control and presented as a fold change relative to the control group.

### Droplet digital PCR

The QX200 Droplet Digital PCR (ddPCR™) system (BioRad) was used for ddPCR analysis. The master mix contained ddPCR supermix (900 nM), primers and probes mix (250 nM), (Supplemental Table 9; IDT Integrated DNA Technologies) for a final 20 μl volume of reaction mix. Droplets were generated by combining 20 µl of the reaction mix with 70 µl of droplet generation oil for probes in the QX200 droplet generator (BioRad). The resulting droplet solution was transferred to a 96 well-PCR plate, the cycling process was carried out and analysed with QuantaSoft software (BioRad).

### Human primary cell culture and transfection

Human Primary Aortic Endothelial Cells (HAoECs) from a 61-year-old female donor (458Z035.1; C-12271; PromoCell) or a 50-year-old male donor (434Z005.1; C-12271; PromoCell) were seeded on 0.2% gelatin-coated plates or ibidi chambers in Endothelial Cell Growth Medium (ECGM; C-22010, PromoCell) supplemented with 1% Penicillin-Streptomycin at 37°C in a humidified atmosphere containing 5% CO2. Cells were used at passage 4 to 6 for experiments. Off-target-plus siRNAs (Smartpool) against human *CNR1* (*siCNR1*) and scrambled siRNA (*siScr*) were obtained from Dharmacon (Supplemental Table 10; Horizon, UK). HAoECs were transfected with 20 nM siRNA utilizing RNAiMax (Invitrogen) and analyzed 24 h or 48 h after transfection.

### Shear stress assays

An ibidi pump system was used to impose laminar (LSS) or oscillatory shear stress (OSS) on confluent monolayers of HAoECs seeded into flow chambers as described.^18,19^ HAoECs were exposed to LSS (10 dyne/cm²) or OSS (3 dyne/cm²) in ECGM for 24 h.

### Monocyte adhesion assay

HAoECs were incubated with CB1 agonist ACEA (1 µM) or vehicle control under LSS at 10 dyne/cm² in ibidi chambers for 24 h. Subsequently, the cells were stimulated with 5 ng/ml TNF-α (R&D) overnight without the application of shear stress. Then, monocytes (1×10^5^ THP-1 /ml) labelled with 0.5 µM calcein (Invitrogen) were added to the flow system. THP-1 and HAoECs were co-cultured under LSS at 37°C for 3 h. Afterwards, the chambers were disconnected from the perfusion system, washed with PBS and imaged with an inverted microscope Leica DMi8 S Platform with the Tilescan function) at 20x magnification. At least 10-15 different random views per condition were selected to quantify the number of adherent THP-1 cells using ImageJ software.

### *In vitro* DiI-LDL uptake

HAoECs were first incubated with CB1 agonist ACEA (1 µM) or vehicle control under LSS for 24 h. Afterwards, 1 µg/ml Dil-LDL was added in medium without supplements and incubated under static condition at 37C° for 1.5 h. Cells were washed, fixed with 2% PFA and permeabilized with 0.1% Triton X-100 for 10 min at room temperature. The chambers were mounted with ibidi Mounting Medium containing DAPI and imaged using an inverted microscope (DMi8 S Platform, Leica) with a 20x objective. The mean fluorescence intensity of Dil-LDL (550nm/564nm) was quantified using LAS V4.3 software (Leica).

In other experiments, AM281 was added 20 min before exposing HAoECs to OSS. The PKA inhibitor KT5720 (1 µM) was added 1 h prior to AM281 treatment. After 24 h of culture in OSS condition, the cells were incubated with 1 µg/ml Dil-LDL for 30 min at 37°C in static condition. The cells were subsequently fixed and permeabilized with 0.1% Triton x-100, followed by incubation with a primary antibody against CAV1 after 1 hour of blocking at room temperature. The next day, the cells were washed and incubated with Alexa Fluor 647 anti-rabbit secondary antibody for 1 hour. The cells were mounted with ibidi Mounting Medium containing DAPI and imaged using an inverted microscope (DMi8 S Platform, Leica) with a 63x objective. Mean Dil-LDL and CAV1 fluorescence intensity were quantified using LAS V4.3 software (Leica) using 5-10 images per condition.

### ROS and nuclear phospho-c-Jun detection

HAoECs were transfected with either 20 nM *CNR1* or scrambled siRNA as described above. After 24 h of transfection, the cells were stimulated with 10 ng/ml TNFα for 30 min. Subsequently, the HAoECs were washed and incubated with 1 µM DHR-123 (Dihydrorhodamine) for 20 min at 37°C to detect ROS production. The cells were then collected after trypsinization, centrifuged, and resuspended in FACS buffer for flow cytometry analysis with a BD FACSCanto II flow cytometer (BD Biosciences). Data were analyzed using FlowJo v10.2 software. For detection of nuclear phospho-c-Jun (Cell signaling), the cells were grown on chamber coverslips (ibidi), treated for 30 min with ACEA, AM281 or vehicle, fixed with 4% PFA and finally immunostained with primary antibody against phospho-c-Jun and secondary with anti-rabbit-AF488. The cells were mounted with ibidi Mounting Medium containing DAPI and images were taken with a fluorescence microscope (DM6000B) connected to a monochrome digital camera (DFC365FX, Leica) equipped with Thunder technology for computational clearing. At least 5 different area views of images per condition were selected for quantification with the Leica Application Suite LAS V4.3 software.

### Cyclic AMP (cAMP) measurement

To measure intracellular cyclic adenosine monophosphate (cAMP) concentrations, HAoECs seeded in 12 well plates were pretreated with 3-isobutyl-1-methylxanthine (0.5 mM, IBMX, Sigma-Aldrich) for 30 min and then stimulated with CB1 antagonist AM281 or CB1 agonist ACEA for 20 min. Forskolin (3 µM, Sigma) was used as a positive control for assessing maximum cAMP levels. Cells were lysed with 0.1 M HCl and intracellular cAMP levels measured with the Cyclic AMP Select ELISA kit (Cayman Chemical). In other experiments, intracellular cAMP was measured in Flp-In TREx-293 (HEK293) cells (Invitrogen) stably transfected with the human *CNR1* cDNA (Missouri S&T cDNA Resource Center) as well as a luciferase-cAMP reporter plasmid (pGloSensor-20F-vector; Promega). F20 Flp-InT-Rex 293 cells were cultured with DMEM supplemented with 10% fetal calf serum and 1% Pen/Strep and passaged using Trypsin-EDTA (0.05%). After incubation with luciferin-EF (2.5 mM, Promega) at room temperature for 1 hour, cells were stimulated with ACEA or AM281 followed by the addition of forskolin (1µM). The luminescence signal as a readout for intracellular cAMP levels was recorded in real time using the Tecan Infinite F200 PRO microplate reader.

### Statistics

Statistical analysis was conducted using GraphPad Prism version 10.0.2 (GraphPad Software, Inc.). All data are presented as mean ± standard error of the mean (SEM). Normality was assessed using either the D’Agostino Pearson omnibus or Shapiro-Wilk normality test within the software. The Mann-Whitney U test was utilized when the data did not adhere to a normal distribution. For comparisons between two groups, a student’s unpaired t-test was applied if the data exhibited a Gaussian distribution (a normal distribution with equal variances) after confirming variances through an F-test. In the context of multiple groups with consideration for a single variable in comparisons, either one-way analysis of variance (ANOVA) with Tukey post hoc test or repeated-measures one-way ANOVA with Tukey post hoc test was used, depending on the experimental design. When two independent factors were involved, a two-way ANOVA with a Bonferroni post hoc test was applied for statistical testing. Outliers were identified using Tukey’s method. P-values less than 0.05 were considered statistically significant.

## SUPPLEMENTAL TABLES

**Supplemental Table 1:** Antibodies used for immunostaining.

| Antigen | Reference | Host | Dilution | Provider |
| --- | --- | --- | --- | --- |
| Caveolin-1 | PA5-17447 | Rabbit | 1:100 | Invitrogen |
| CD31 (PECAM-1) | 553370 | Rat | 1:50 | BD |
| CD54 (ICAM1) | 553250 | Hamster | 1:100 | BD |
| CD68 | MCA1957GA | Rat | 1:400 | Bio-Rad |
| CD106 (VCAM1) | 553329 | Rat | 1:100 | BD |
| CD144 (VE-Cadherin) | 555289 | Rat | 1:100 | BD |
| GPIHBP1 | PA5-98598 | Rabbit | 1:100 | Invitrogen |
| Hoechst | H1399 |  | 1:1000 | Invitrogen |
| LPL | AF7197-SP | Goat | 1:100 | R&D system |
| Phospho-c-Jun | 3270T | Rabbit | 1:100 | Cell Signaling |
| vWF (Von Willebrand Factor) | Ab11713 | Sheep | 1:300 | Abcam |

Blocking buffer: 6 ml PBS, 600 µl 10 % BSA (1 %), 3 drops horse serum (S-2000, Vector Laboratories). Antigen Retrieval buffer: 630 ml Aqua dest, 12,6 ml Solution A (21,01 g Citric Acid, 1 L Aqua dest), 57,4 ml Solution B (29,4 g Sodium Citrate, 1 L Aqua dest), 350 µl Tween20.

**Supplemental Table 2:** Secondary antibodies.

| Antigen | Source | Conjugation | Dilution | Reference | Provider |
| --- | --- | --- | --- | --- | --- |
| Anti-goat | Donkey | AlexaFluor594 | 1:100 | 705-585-003 | JIR |
| Anti-hamster | Goat | Cy3 | 1:300 | 127-165-160 | JIR |
| Anti-rabbit IgG | Donkey | AlexaFluor647 | 1:600 | 711-605-152 | JIR |
| Anti-rabbit IgG | Donkey | Cy3 | 1:300 | 711-165-152 | JIR |
| Anti-rat | Donkey | AlexaFluor488 | 1:300 | A21208 | Invitrogen |
| Anti-rat | Donkey | Cy3 | 1:300 | 712-165-153 | JIR |
| Anti-sheep | Donkey | DyLight® 488 | 1:300 | ab96939 | Abcam |
| Anti-sheep IgG | Donkey | Cy5 | 1:600 | 713-175-147 | JIR |
| Anti-sheep IgG | Donkey | Cy3 | 1:600 | 713-165-003 | JIR |

**Supplemental Table 3:**
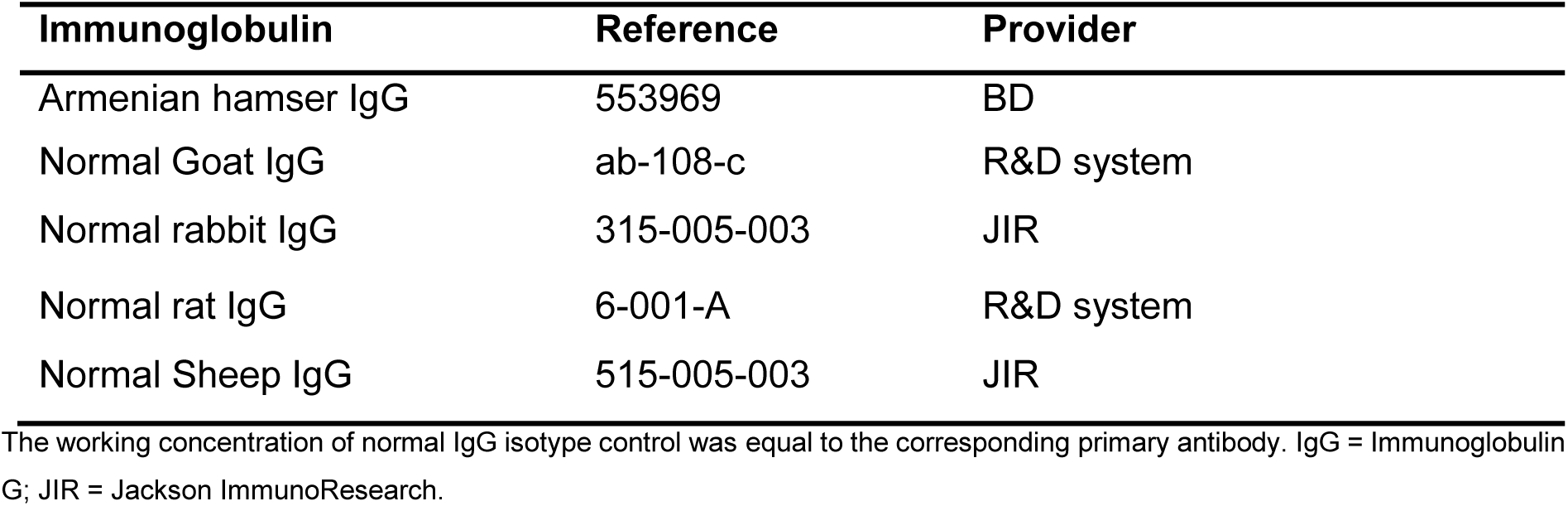
Isotype controls.

**Supplemental Table 4:** Antibodies used for TPLSM.

| **Antigen** | **Reference** | **Host** | **Dilution** | **Provider** |
| --- | --- | --- | --- | --- |
| CD31/eFluor450 (PECAM-1) | 48-0311-82 | Rat | 1:100 | eBioscience™ |
| Dil-LDL | 770230-9 | human | 1:10 | KalenBiomedica |

**Supplemental Table 5:** Murine flow cytometry antibodies.

| Antigen | Conjugation | Dilution | Reference | Provider |
| --- | --- | --- | --- | --- |
| CD11b | PerCP | 1:500 | 101230 | Biolegend |
| CD16/32 | purified | 1:1000 | 553142 | BD |
| CD31 | PE-Cy7 | 1:100 | 102417 | Biolegend |
| CD36 | APC | 1:1000 | Ab133625 | Abcam |
| CD45 | Alex eFluoro 780 | 1:400 | 47-0451-82 | Invitrogen |
| CD45.2 | FITC | 1:500 | 553772 | BD |
| CD54 (ICAM-1) | APC | 1:500 | 116119 | Biolegend |
| CD106 (VCAM-1) | PerCP | 1:500 | 105715 | Biolegend |
| CD107a | BV421 | 1:400 | 121617 | Biolegend |
| CD115 | APC | 1:500 | 17-115-282 | eBioscience |
| Live/dead | Zombie Green | 1:800 | 77476 | Biolegend |
| LOX1 | AF647 | 1:100 | FAB1564R | R&D system |
| Ly6C | PE- Cy7 | 1:500 | 560593 | BD |
| Ly6G | APC-Cy7 | 1:500 | 127623 | Biolegend |
| SRA1 | FITC | 1:500 | ab151707 | Abcam |
| SRB1 | FITC | 1:500 | NB400-104F | NOVUS |

APC = Allophycocyanin; BV = Brilliant violet; Cy = Cyanine; FC = Flow Cytometry; FITC = Fluorescein isothiocyanate; IF=Immunofluorescence; PB = Pacific Blue; PE = Phycoerythrin; PerCP = Peridinin chlorophyll.

**Supplemental Table 6:**
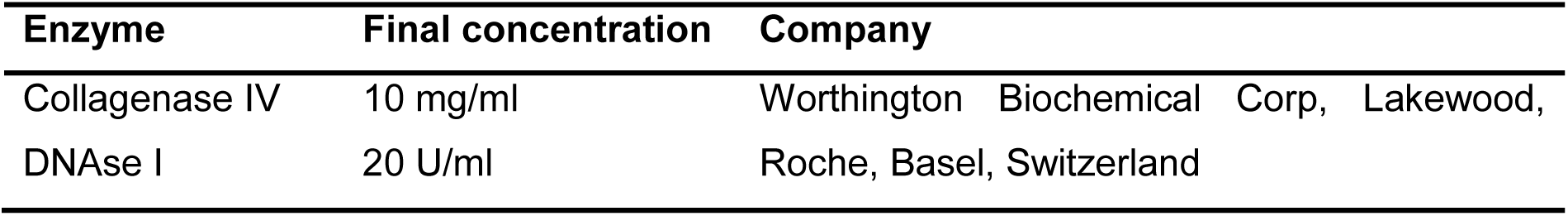
Enzymes for aorta digestion.

**Supplemental Table 7:**
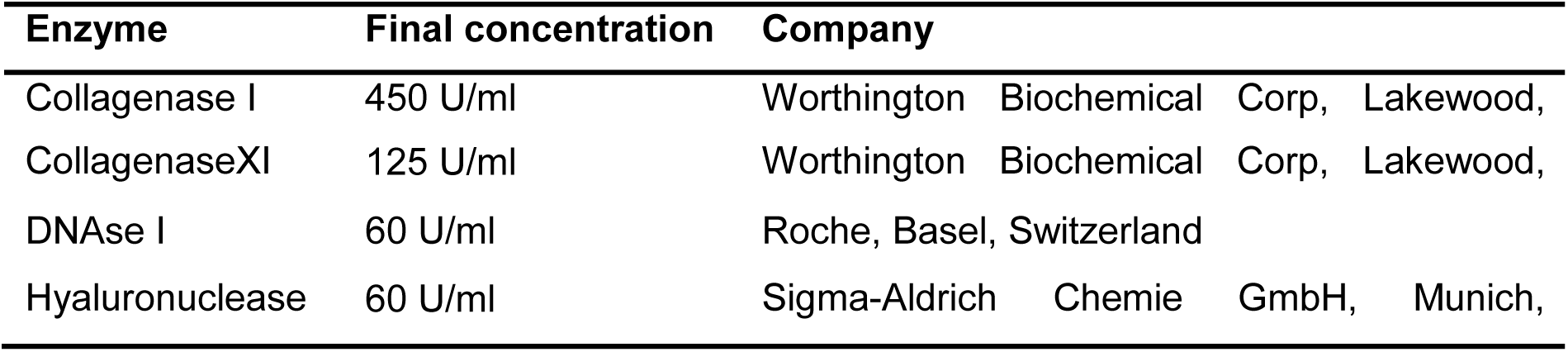
Enzymes for BAT digestion.

**Supplemental Table 8:**
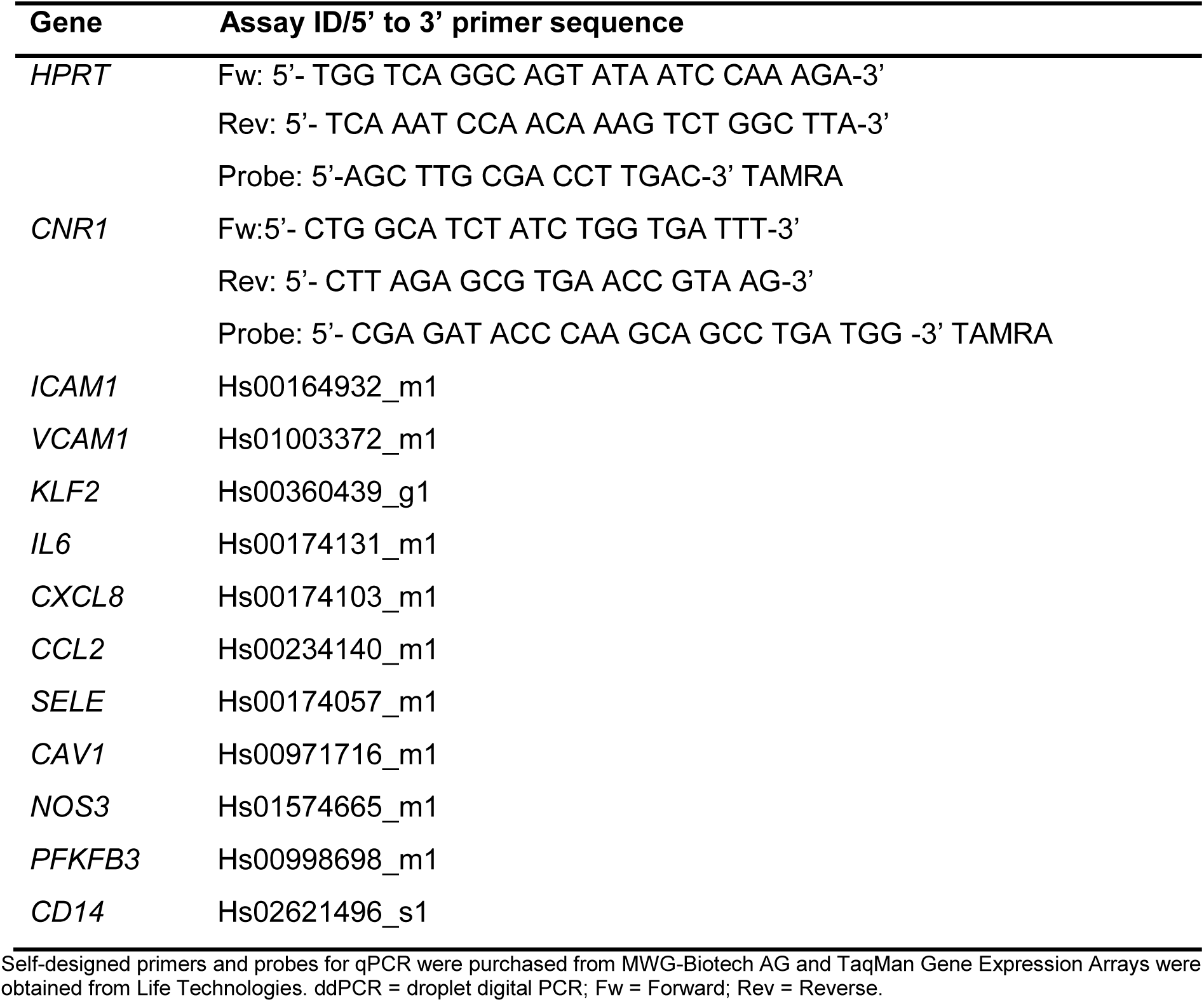
Primers for qPCR analysis (Human)

**Supplemental Table 9:**
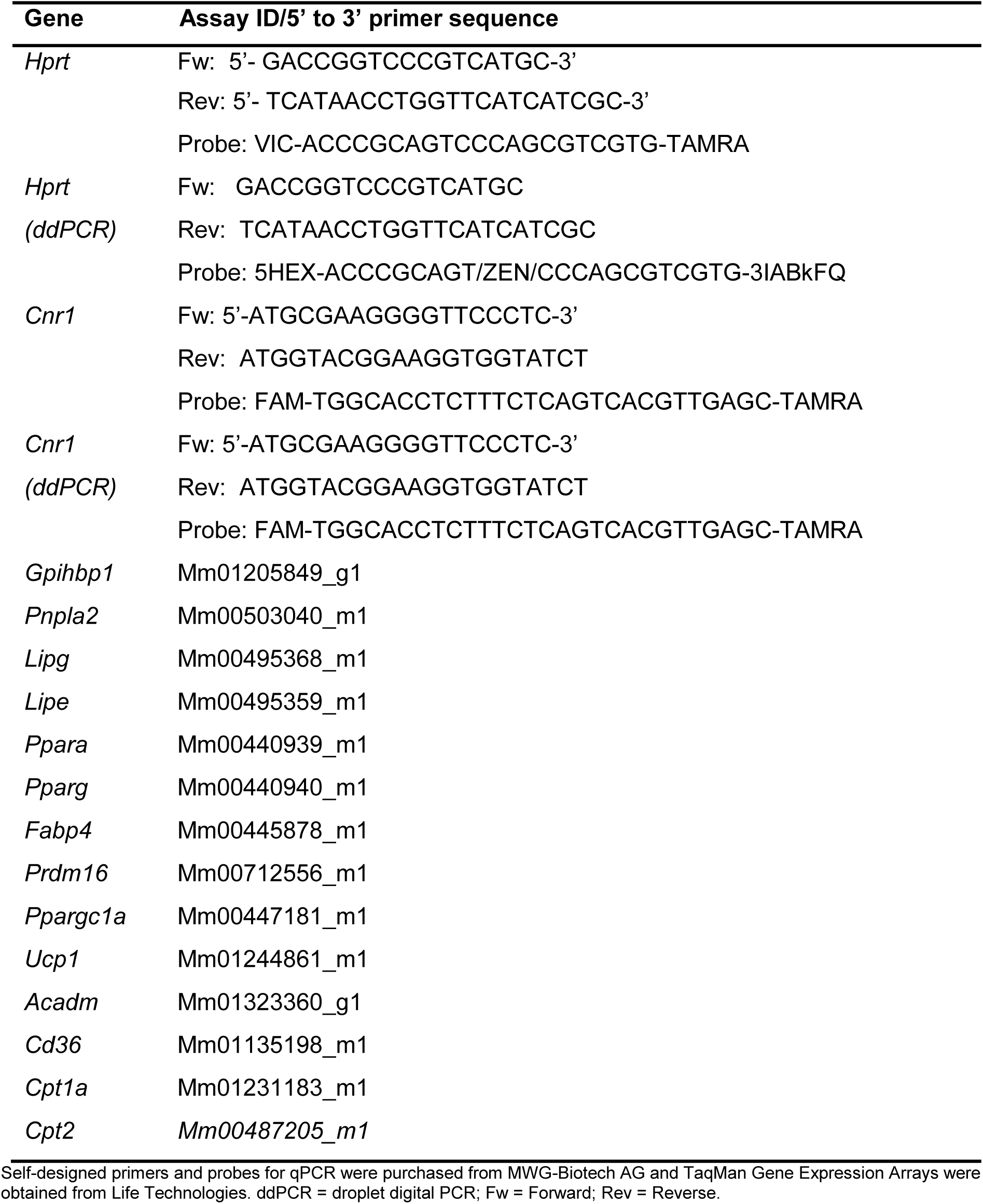
Primers for qPCR analysis (Murine)

**Supplemental Table 10:**
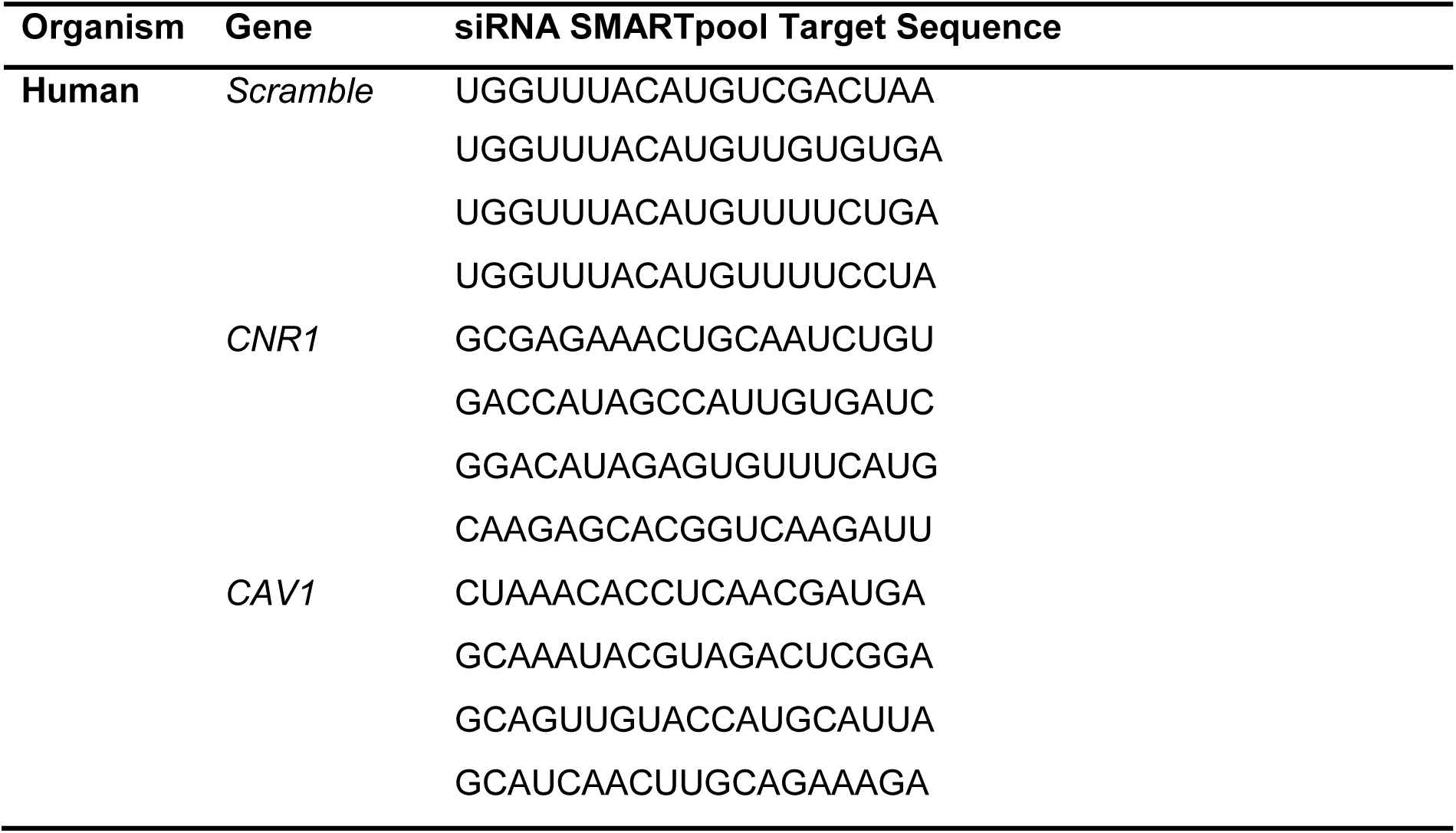
siRNA SMARTpool Target Sequence.

